# A critical period of translational control during brain development at codon resolution

**DOI:** 10.1101/2021.06.23.449626

**Authors:** Dermot Harnett, Mateusz C. Ambrozkiewicz, Ulrike Zinnall, Alexandra Rusanova, Ekaterina Borisova, Rike Dannenberg, Koshi Imami, Agnieszka Münster-Wandowski, Beatrix Fauler, Thorsten Mielke, Matthias Selbach, Markus Landthaler, Christian M.T. Spahn, Victor Tarabykin, Uwe Ohler, Matthew L. Kraushar

**Author notes:** These authors contributed equally (Dermot Harnett, Mateusz C. Ambrozkiewicz). Senior author (Victor Tarabykin, Uwe Ohler, Matthew L. Kraushar). Correspondence (D.H.); (M.C.A.); (M.L.K.).

## Abstract

Translation modulates the timing and amplification of gene expression after transcription. Brain development requires uniquely complex gene expression patterns, but large-scale measurements of translation directly in the prenatal brain are lacking. We measure the reactants, synthesis, and products of translation spanning mouse neocortex neurogenesis, and discover a transient window of dynamic regulation at mid-gestation. Timed translation upregulation of chromatin binding proteins like Satb2, which is essential for neuronal subtype differentiation, restricts protein expression in neuronal lineages despite broad transcriptional priming in progenitors. In contrast, translation downregulation of ribosomal proteins sharply decreases ribosome number, coinciding with a major shift in protein synthesis dynamics at mid-gestation. Changing levels of eIF4EBP1, a direct inhibitor of ribosomal protein translation, are concurrent with ribosome downregulation and controls Satb2 fate acquisition during neuronal differentiation. Thus, the refinement of transcriptional programs by translation is central to the molecular logic of brain development. Modeling of the developmental neocortex translatome is provided as an open-source searchable resource: https://shiny.mdc-berlin.de/cortexomics/.

## Introduction

Changes in translation activity can lead to significant discrepancies between mRNA and protein for the same gene, and are a hallmark of many dynamic cellular transition states ^1^. Dynamic cellular transitions are uniquely complex during brain development from neural stem cells, which must deploy highly sophisticated gene expression programs ^2,3^. In evolutionarily advanced brain regions like the neocortex, a cell’s transcriptional signature alone appears insufficient to account for the enormous cellular diversity, with recent single-cell RNA sequencing (scRNA-seq) analyses supporting this idea ^4–7^. While transcriptional profiles define broad classes of neurons and non-neuronal cells, a striking conclusion from these studies was the degree of homogeneity in mRNA pools across neuronal lineages during stem cell differentiation ^4^, and between distinct neuronal circuits postnatally ^7^. Considering whether neuronal differentiation in the neocortex utilizes a more “generic” transcriptome ^4^ has led the field to ask recently whether neuronal identity is a stochastic rather than deterministic process ^8^. Do “progenitors play dice” ^9^ while deciding their neuronal fate? Thus, the blueprint of gene expression in evolutionarily advanced brain regions is likely a multilayered, progressive refinement – including and beyond transcription ^3^. Neocortex development may thus represent a particularly dynamic cellular system of translational control ^10,11^.

Direct measurements of protein synthesis would provide a clearer picture of functional gene expression in the developing brain; however, a large-scale high-resolution analysis of mRNA translation during neurogenesis has lagged behind transcriptome analysis, in part due to current technical limitations in protein measurement. Recent work suggests that targeted and selective protein synthesis refines the output of gene expression in brain development ^5,12–16^. Importantly, abnormal ribosome levels and disrupted translation was found recently to be a mid-gestation etiology of neurodevelopmental disorders ^17^. However, how ribosomes decode mRNA in the transcriptome-to-proteome transition during developmental neurogenesis remains unknown.

To circumvent these challenges and measure the temporal dynamics of the reactants, synthesis, and products of mRNA translation during brain development, we performed sequencing of ribosome-protected mRNA fragments (Ribo-seq; ribosome profiling) ^18^ in parallel with RNA-seq, tRNA qPCR array, and mass spectrometry across five stages of mouse neocortex neurogenesis. By capturing ribosome-mRNA interactions at codon-level resolution, we find that ~ 18 % of mRNAs change translation efficiency in the progressive specification of neural stem cells to post-mitotic neurons, with a transient peak window of dynamic translation at mid-gestation.

Divergent cellular pathways are impacted by translation upregulation vs. downregulation during neurogenesis. Chromatin binding proteins like *Satb2*, essential for the differentiation of neuronal subtypes ^19^, are the most translationally upregulated mRNAs. We find *Satb2* mRNA is transcribed unexpectedly broadly in neuronal lineages, but achieves restricted neuronal subtype-specific protein expression by timed translation. In contrast, ribosomal proteins are the most translationally downregulated, coinciding with dynamic expression of eIF4EBP1, a translational repressor targeting these transcripts directly ^20,21^. An acute decrease in ribosome number coincides with widespread changes in global translation kinetics at mid-gestation – a critical period for neurodevelopmental pathology ^17^. Finally, *in utero* knockdown of eIF4EBP1 in neural progenitors to disrupt the balance of translation during this window leads to decreased specification of the Satb2 neuronal lineage and migration arrest.

Thus, by mapping the quantitative landscape of the transcriptome-to-proteome transition in the neocortex, we find that protein synthesis is a powerful and widespread layer of gene expression regulation that shifts kinetics and impacts neuronal specification during development. We provide the developmental neocortex translatome as an open-source searchable web resource: https://shiny.mdc-berlin.de/cortexomics/.

## Results

### Deep sequencing of translation reveals a spike in regulation during neurodevelopment

We focused our study on the mammalian neocortex, an evolutionarily advanced and dynamic developmental system with a tightly timed sequence of neurogenesis ^2,22^ (**Fig. 1a**). At embryonic day 12.5 (E12.5) this predominantly stem cell tissue gives birth to its first neurons. Neurons born early at E12.5 form distinct connections and control different functions than those born later at E15.5. By postnatal day 0 (P0), neurogenesis is largely complete. The timed sequence of gene expression is essential to specify neuronal fate from the stem cell pool.

**Fig. 1.**
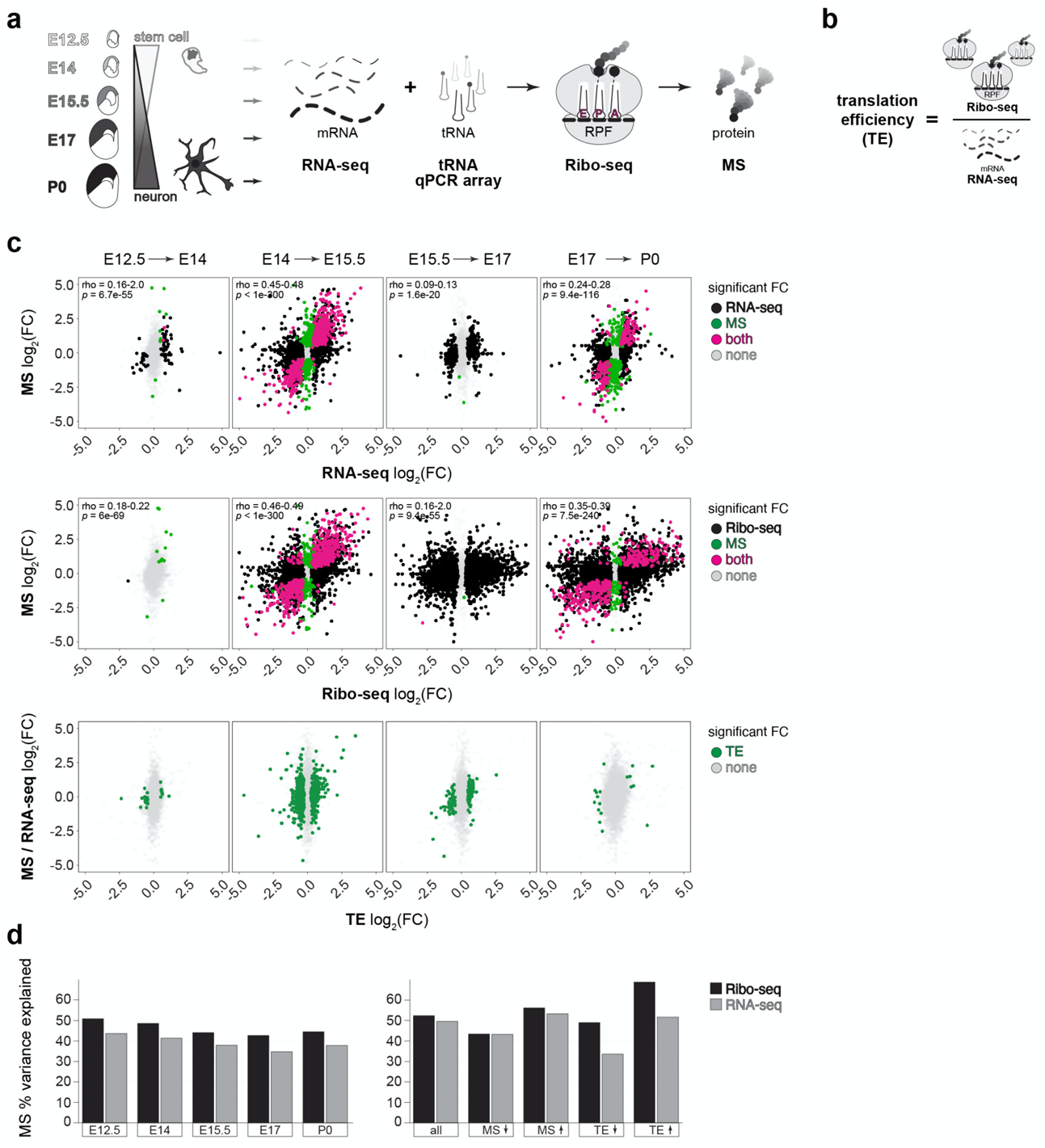
A transient spike in translation regulation occurs at mid-neurogenesis during prenatal development. **a**, Neural stem cell differentiation in the brain’s neocortex, analyzed by RNA-seq, Ribo-seq, tRNA qPCR array, and MS at embryonic (E12.5-E17) and postnatal (P0) stages. **b**, Schematic of translation efficiency (TE). **c**, Sequential fold changes in post-transcriptional gene expression between adjacent stages, comparing mRNA vs. protein (top), mRNA translation vs. protein (middle), and calculated translation efficiency (bottom). Significance assessed at ≥ 1.25 FC, *p* < 0.05. **d**, The percent variance in MS explained by RNA-seq or Ribo-seq at each developmental stage, and for subgroups with MS and translation efficiency changes. See also **Extended Data Figs. 1–2 and Supplementary Tables 1-2**.

We designed a strategy to analyze the major reactants, synthesis, and products of mRNA translation across five stages encompassing neocortex neurogenesis (**Fig. 1a**), including Ribo-seq measurement of ribosome-mRNA interactions, in parallel with RNA-seq, tRNA qPCR array, and mass spectrometry. Ribo-seq measures 80S ribosomes bound to the open reading frame of mRNA – a quantitative indicator of active protein synthesis at codon-level resolution ^18^. Optimizations for analysis of neocortex ribosomes *ex vivo* circumvented the requirement for pharmacological ribosome stalling with cycloheximide ^15^, which introduces ribosome footprint redistribution artifacts ^23^, and enabled efficient nuclease digestion to generate high fidelity ribosome-protected mRNA fragments (RPFs) (**Extended Data Fig. 1**). We obtained mRNA transcripts per-million (TPM) and RPF densities for 22,373 genes (**Extended Data Fig. 2a, Supplementary Table 1**). Reproducibility of both mRNA and RPF measurements permitted reliable calculation of mRNA translation efficiency (**Extended Data Fig. 2b-e**), which represents the ratio between ribosome binding to an mRNA’s coding sequence and the mRNA’s level overall (**Fig. 1b**). As a quality control, we focused further analysis on coding sequences with 32 or more Ribo-seq footprints in at least one stage as per ^24^, which resulted in a set of 12,228 translated GENCODE-annotated transcripts.

**Fig. 2.**
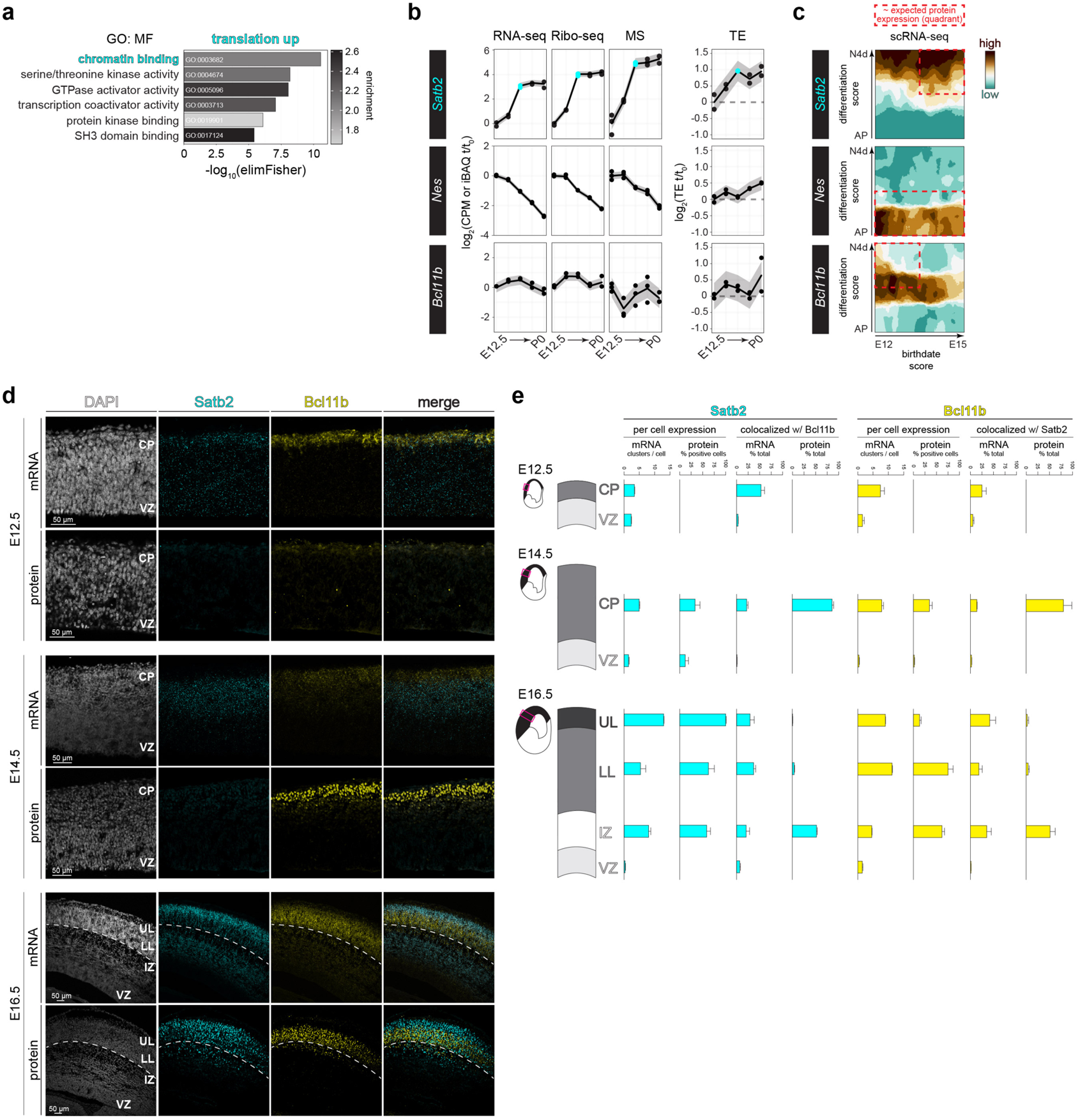
Translation upregulation of *Satb2* leads to divergent spatiotemporal mRNA and protein expression. **a**, Gene ontology (GO, molecular function) analysis of translationally up-regulated (TE up) mRNAs. **b**, The median trajectory of *Satb2, Nes,* and *Bcl11b* gene expression measured by RNA-seq, Ribo-seq, MS, and translation efficiency. The E15.5 timepoint is highlighted for Satb2 **c**, *Satb2, Nes,* and *Bcl11b* expression in scRNA-seq data tracking differentiating neocortex cells at 1, 24, or 96 hours after birth (y-axis), at birthdates E12, E13, E14, or E15 (x-axis) (Telley et al., 2019). Expected distribution of protein expression (DeBoer et al., 2013) is outlined. **d**, Neocortex coronal sections at E12.5, E14.5, and E16.5 analyzed for *Satb2* and *Bcl11b* mRNA by fluorescence *in situ* hybridization, and protein by immunohistochemistry. Deep border of the cortical plate is demarcated at E16.5 (dotted line). DAPI (nuclei). Ventricular zone (VZ), cortical plate (CP), lower layers (LL), upper layers (UL). **e**, Quantification of (d). Mean ± SEM. See also **Extended Data Fig. 3a-b and Supplementary Table 3**.

To detect translation-specific gene expression regulation, we first calculated fold changes between sequential time points in mRNA or RPF vs. protein (**Fig. 1c, Supplementary Table 2**). While gene expression overall is quite stable between E12.5 and E14, a burst of regulation occurs at E15.5 at both transcriptional and translational levels, with a significant impact on the proteome. However, robust RPF fold changes persist until P0, with mRNA changes less pronounced. Calculation of translation efficiency highlighted a transient window of robust regulation at E15.5, coinciding with the major transition in neuronal fate specification. Translation efficiency upregulation was found to occur in 1,129 genes and downregulation in 1,131 genes. A further 2,253 genes change in steady state mRNA only, without any significant translation efficiency change. Thus, we estimate ~ 18 % of the transcriptome is dynamically translated across neocortex neurogenesis, with an acute inflection point during the mid-neurogenesis transition at E15.5.

Our Ribo-seq data shows a higher correlation with protein level changes than RNA-seq data, (**Fig. 1c**). We decomposed technical and systematic variation in protein levels, and estimated proportions explained by RNA-seq vs. Ribo-seq ^25^ (**Fig. 1d and Extended Data Fig. 2f**). A majority of protein level variance is accounted for by RNAseq, in agreement with prior observations ^25,26^. However, Ribo-seq consistently explains a higher fraction of protein variation than RNA-seq at each developmental stage, especially for proteins with increasing levels – concordant with Ribo-seq being a direct measure of protein synthesis. The protein level predictivity of Ribo-seq was particularly pronounced for genes with changing translation efficiency.

Thus, our data enable detection of regulation that impacts the protein output of gene expression, which includes a transient window of robust translation control at E15.5.

### Translation upregulation of chromatin binding proteins like Satb2 establishes neuronal fate

We first focused on the cohort of genes that are translation efficiency upregulated across neurogenesis after E12.5, to identify essential neurodevelopmental proteins with dynamic translation. Gene ontology analysis demonstrated chromatin binding proteins are particularly subject to translation upregulation (**Fig. 2a**). Chromatin binding proteins like transcription factors have a powerful influence on the neuronal fate of stem cells, which is tightly coordinated in developmental time. *Early*-born post-mitotic neurons ultimately express transcription factors like Bcl11b, which drives them to connect *sub*-cortically ^27^. In contrast, *late*-born post-mitotic neurons after E15.5 ultimately express transcription factors like Satb2, which drives them to connect *intra*-cortically ^19,28^. How proteins like transcription factors achieve neuronal subtype and temporally restricted expression is a critical unresolved question.

Among the most translationally upregulated neurodevelopmental proteins discovered in our data is the essential, late-born upper layer neuron transcription factor Satb2 (**Fig. 2b**). We assessed the trajectory of Satb2 synthesis in our RNA-seq, Ribo-seq, and mass spec data along with calculated translation efficiency; in comparison to the intermediate filament protein Nes expressed by neural stem cells ^29^, and early-stage transcription factor Bcl11b expressed in neurons positioned adjacent to the later Satb2 lineage. As expected, Nes demonstrates predominantly transcriptionally driven expression downregulation, as the neural stem cell pool is depleted by neuronal differentiation ^30^. Bcl11b is expressed in the early-born lineage with high concordance between RNA-seq and Ribo-seq, and with low fluctuations in translation efficiency. In contrast, fold changes in Satb2 Ribo-seq and MS signal are in excess of the RNA-seq, with 2-fold translation efficiency upregulation reaching a plateau at E15.5. These data suggest that *Satb2* expression is amplified by translation.

To begin testing the hypothesis that *Satb2* mRNA undergoes translation regulation, we first examined the cellular distribution of *Satb2* mRNA in scRNA-seq neuronal lineage-tracing data ^4^. Surprisingly, we found that *Satb2* mRNA is robustly expressed in differentiated neurons of both the early- *and* late-born lineages (**Fig. 2c**) – an apparent discrepancy with previous findings for Satb2 protein ^19,28^. Thus, transcription of this upper layer program may occur in neuronal lineages that include lower layers, and outside of the expected protein distribution.

To directly visualize the spatiotemporal expression of *Satb2* mRNA and protein, we performed fluorescence *in situ* hybridization and immunohistochemistry in neocortical coronal sections (**Fig. 2d**), with probe and antibody specificity confirmed in *Satb2*^-/-^ brains (**Extended Data Fig. 3a**), and signal quantified per cell (**Fig. 2e and Extended Data Fig. 3b, Supplementary Table 3**). At the onset of neurogenesis E12.5, initial scattered, weak Bcl11b protein signal is congruent with its mRNA signal in post-mitotic neurons. Satb2 protein is undetectable; however, we observed robust *Satb2* mRNA signal throughout the neocortex, from the ventricular zone in multipotent progenitors and throughout the nascent cortical plate in early-born post-mitotic neurons. In neurons differentiating in the cortical plate, almost half of all *Satb2* mRNA clusters colocalize with *Bcl11b* mRNA, which rarely occurs in the stem cell niche of the ventricular zone.

**Fig. 3.**
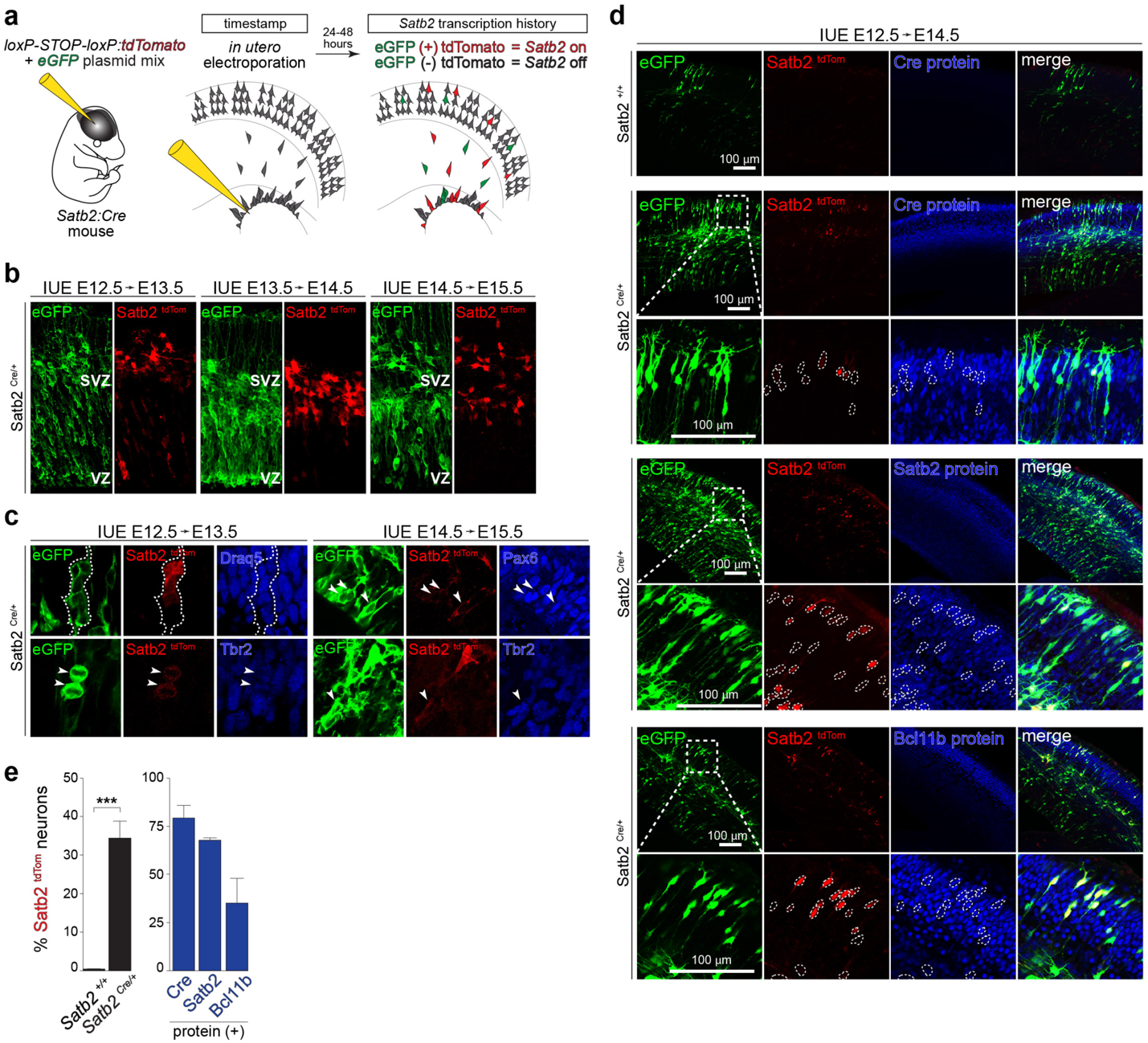
*Satb2* transcription is broad across neuronal lineages with more restricted translation. **a**, Schematic of the experimental approach. **b**, *Satb2* transcription activity visualized by Cre-driven (*Satb2^Cre/+^*) tdTomato expression, with reporter *in utero* electroporation (IUE) at E12.5, E13.5, or E14.5 and imaged after 24 hours. Co-electroporation of an eGFP plasmid labels all transfected cells. Ventricular zone (VZ), subventricular zone (SVZ). **c**, *Satb2^tdTom^* co-immunolabeling with Pax6 (apical progenitors), Tbr2 (intermediate progenitors), and Draq5 (nuclei). **d**, *Satb2^tdTom^* expression at E12.5-E14.5, with co-immunolabeling for neuronal fate determinant proteins Satb2 and Bcl11b, among all electroporated cells (eGFP). Negative control is the absence of *Cre* (Satb2^+/+^). **e**, Quantification of (d) for the expression of Cre, Satb2, and Bcl11b proteins in cells transcribing *Satb2* mRNA (*Satb2 ^tdTom^*). Mean ± SD, unpaired t-test, \*\*\**p* < 0.0001. See also **Extended Data Fig. 3c and Supplementary Table 3**.

Weak Satb2 protein expression is first detected at E14.5, in contrast to strong Bcl11b protein now appearing in post-mitotic neurons. Only by E16.5 is Satb2 protein expression robust. Satb2 mRNA and protein are broadly expressed across upper layers, lower layers, and the intermediate zone by E16.5. However, neurons having migrated to their ultimate position in upper layers almost exclusively express Satb2 rather than Bcl11b protein, in contrast to regions like the intermediate zone where neurons continue to migrate.

Taken together, *Satb2* mRNA and protein expression are divergent in developmental time and space. This divergence includes broad, early *Satb2* mRNA expression in multipotent progenitors despite Satb2 protein ultimately restricted to upper layer post-mitotic neurons later in development. Furthermore, while the distribution and colocalization of mRNA for *Bcl11b* and *Satb2* neuronal programs remains broad and overlapping in post-mitotic neurons at E16.5, corresponding protein expression is more exclusive, with the intermediate zone a transitory region where specification at both the mRNA and protein levels are still lacking distinction. Thus, our bioinformatics analysis identifies *Satb2* as a translationally upregulated mRNA, for which we observe incongruent spatiotemporal mRNA-protein expression *in situ*.

### Translation establishes the balance of neuronal fates after broad transcription

Given the unexpected finding of *Satb2* mRNA in early-born neural stem cells and overlap with the *Bcl11b* neuronal lineage, we next sought to monitor transcriptional activation of the *Satb2* locus. We employed a fate mapping approach with the *Satb2*^Cre/+^ mouse line ^31^. A Cre expression cassette is located in place of exon 2 at the *Satb2* locus, which allows for timed *in utero* electroporation of Cre-inducible reporters like *loxP-STOP-loxP-tdTomato* that clonally labels cells with tdTomato that have a history of *Satb2* transcription (*Satb2^tdTom^*) (**Fig. 3a**). Co-electroporation with an *eGFP* plasmid serves as a generic label for all transfected cells.

Remarkably, we detected *Satb2^tdTom^* cells in the ventricular zone as early as E12.5 forming clusters resembling clones or undergoing mitotic divisions (**Fig. 3b**), and express neural progenitor markers like Pax6 (apical progenitors) or Tbr2 (intermediate progenitors) (**Fig. 3c**). *Satb2* transcription was observed for progenitors in the neocortex, but not in adjacent brain regions (**Extended Data Fig. 3c**). Thus, *Satb2* transcriptional priming occurs in early-born neocortex neural stem cells, indicating broad transcription of a protein expressed in a restricted neuronal lineage appearing later.

The balance of the Bcl11b vs. Satb2 lineages is essential for normal neocortex development and function. Satb2 directly suppresses the *Bcl11b* genomic enhancer, and loss of Satb2 engenders ectopic expression of *Bcl11b* in upper layer neurons, leading to abnormal connectivity ^19^. Therefore, we next investigated the expression of Bcl11b and Satb2 protein in cells that transcribe *Satb2* mRNA (**Fig. 3d-e, Supplementary Table 3**). Among cells transcribing *Satb2* mRNA, ~70% express Satb2 protein and ~30% express Bcl11b protein. Taken together, this observation indicates that despite unexpectedly broad and early transcription of the neuronal fate gene *Satb2*, translation of Satb2 protein restricts its expression to a late-born neuronal subtype, and maintains the balance of alternative neuronal fates.

### Translation downregulation decreases ribosome levels acutely at mid-neurogenesis E15.5

We next focused on genes that are translationally downregulated across neurogenesis after E12.5. Gene ontology analysis highlighted structural constituents of the ribosome, predominantly ribosomal proteins, as strongly downregulated by translation (**Fig. 4a**). We calculated the developmental expression trajectory of all 79 ribosomal proteins in the large and small subunits by RNA-seq, Ribo-seq, mass spec, and translation efficiency (**Fig. 4b**). Results showed downregulation of nearly all ribosomal proteins at the Ribo-seq and MS level occurs acutely at E15.5, in advance of changes measured by RNA-seq, and reflecting translation downregulation until mid-neurogenesis. Decreasing ribosome levels by downregulation of ribosomal protein translation likely represents the coordinated regulation of this specific gene family, rather than a simple translation feedback loop, since numerous genes in other families undergo translation upregulation concurrently, such as chromatin binding proteins.

**Fig. 4.**
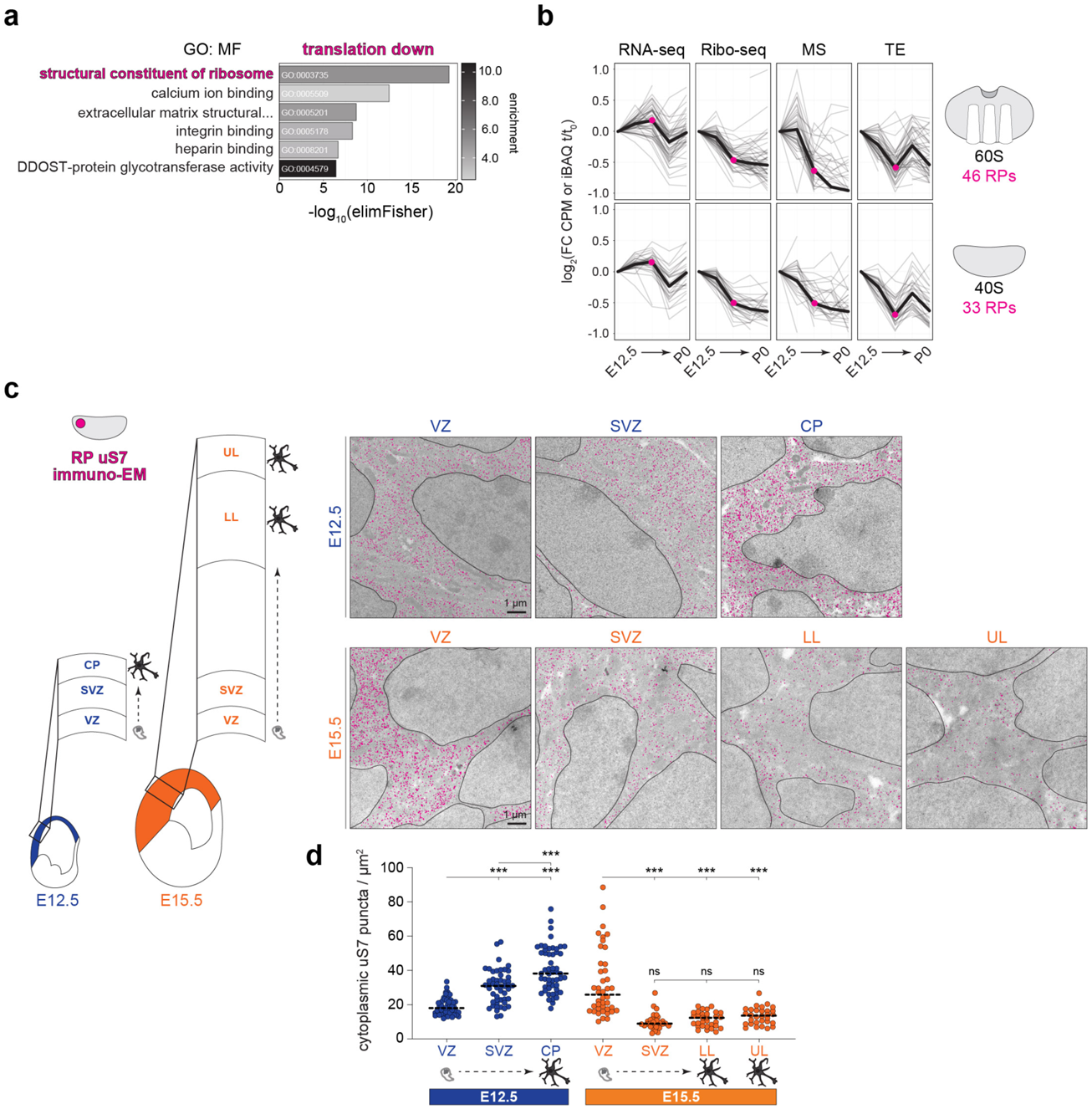
Translation downregulation decreases ribosome levels acutely at mid-neurogenesis. **a**, Gene ontology (GO, molecular function) analysis of translationally down-regulated (TE down) mRNAs. **b**, The expression trajectories (grey) of all 79 ribosomal protein coding mRNAs in the large (*Rpl*) and small (*Rps*) subunits from E12.5 (t_0_) to subsequent stages (t), measured by RNA-seq, Ribo-seq, MS, and calculated translation efficiency. Median trajectories are shown (black). **c**, Immuno-electron microscopy labeling ribosomal protein uS7 (magenta) in the E12.5 and E15.5 neocortex neural stem cells and neurons, with **d**, quantification for ribosomes per cytoplasmic area. Mean shown (line), Welch’s ANOVA and Dunnett’s *post hoc* test, \*\*\**p* < 0.001. Neural stem cells are located in the ventricular zone (VZ) and sub-ventricular zone (SVZ); post-mitotic neurons are located in the cortical plate (CP), lower layers (LL), upper layers (UL). Nuclei are outlined. See also **Extended Data Fig. 4 and Supplementary Table 3**.

To detect changing ribosome numbers sub-cellularly at high resolution, we performed immuno-electron microscopy analysis labeling ribosomal protein uS7 at E12.5 and E15.5 in the neocortex (**Fig. 4c-d and Extended Data Fig. 4, Supplementary Table 3**). A striking decrease in ribosome number was observed in differentiating neurons from early to late stages. Ribosomes are abundant in cortical plate neurons at E12.5, but scarce in both upper and lower layer neurons of the cortical plate at E15.5. Notably, a progressive increase in ribosome number was observed as newly born neurons traverse away from the ventricular zone into the cortical plate at E12.5; while at E15.5, ribosome numbers decrease precipitously outside the ventricular zone, with few ribosomes measured in sub-ventricular zone progenitors. Thus, ribosome number is temporally enforced by translation at mid-gestation. As ribosome abundance is a powerful determinant of translation kinetics and selectivity ^32,33^, global shifts in translation activity may occur at mid-neurogenesis.

### Ribosome density at the start codon and within the CDS are developmentally dynamic

We next examined global translation activity during neocortex development by determining ribosome-mRNA interactions per-codon across all coding sequences. Ribosome position aligned to codons in the P-site demonstrated the characteristic 3-nucleotide periodicity in Ribo-seq metagene plots (**Fig. 5a**). We found ribosome occupancy surrounding the start codon increases sharply at E15.5, with progressive increases per stage until P0, while stop codon occupancy demonstrates the opposite trend and occurs independent of start codon changes (**Supplementary Table 4**). We applied RiboDiPA ^34^, a linear modeling framework designed for positional analysis of Ribo-seq signal, to pinpoint the ~ 5-fold ribosome occupancy changes to the 4 codon bin surrounding the start and stop (**Fig. 5b**).

**Fig. 5.**
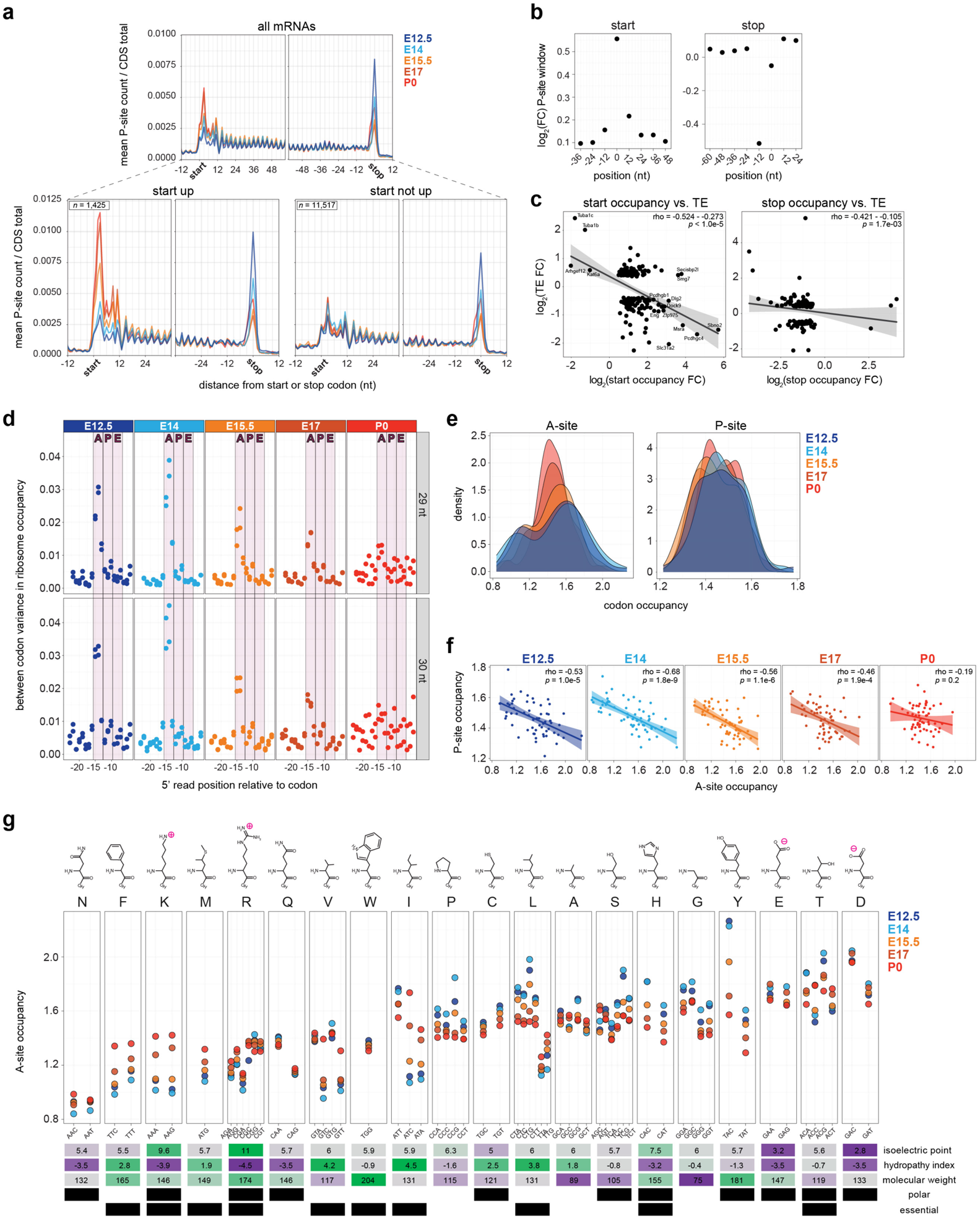
Ribosome density at the start codon and in the CDS shifts sharply at mid-neurogenesis. **a**, Ribosome occupancy metagene plot including all mRNAs (top) surrounding the start (left) and stop (right) codons at five stages. Separation of mRNAs by changing or unchanged start codon occupancy (bottom). **b**, Position specific fold changes in ribosome P-site counts surrounding the start and stop codons. **c**, Start (left) and stop (right) codon occupancy vs. TE fold change per gene. **d**, Between codon variance in ribosome occupancy of A-, P-, and E-sites at each stage. Calculation with both 29 nt (top) and 30 nt (bottom) RPF fragments shown. **e**, Distribution of per-codon A- and P-site occupancy at each stage. **f**, Correlation between A- and P-site occupancy per codon. **g**, Ribosome A-site occupancy for each amino acid with corresponding synonymous codons at each stage. See also **Extended Data Figs. 5–6, Supplementary Table 4**.

Increased ribosome occupancy of the first four codons over time could represent a narrowing bottleneck in the transition from initiation to elongation, or signify increasingly robust initiation of target mRNAs. We correlated fold changes in start codon occupancy with translation efficiency and found an inverse relationship, suggesting that early elongation events progressively slow over time for a large cohort of proteins (**Fig. 5c**). Translation of the N-terminus may become increasingly rate-limiting during synthesis Sbno2 and Pcdhgc4, in contrast to Tuba1c and Tuba1b representing more processive translation during development. Thus, as ribosome levels decline at E15.5 to P0, translation at the 5’-end of coding sequences occurs more slowly.

We next investigated distinct positions where variations in ribosome density take place ^35^ (**Fig. 5d**; see **Methods**). A narrow region consistent with the ribosomal A-site accounts for most of the codon-specific variation in ribosome occupancy. Variation in A-site occupancy was most pronounced at E12.5-E14, with an acute decrease at E15.5-E17, and low variation by P0. Analysis of ribosome dwell time per codon – a measure of the codon-specific speed of translation ^36^ – demonstrated early developmental “fast” or “slow” kinetics in the bimodal distribution of codon dwell times in the A-site (**Fig. 5e and Extended Data Fig. 5a-b; Supplementary Table 4**). At E15.5, codon dwell times begin to equalize, progressively reaching a unimodal distribution by P0. Furthermore, ribosome density occupying A-site codons negatively correlates with P-site density in the embryonic period, but no correlation was measured after birth at P0 (**Fig. 5f**). Thus, the A-site codon in particular influences ribosome dwell time, which is a barrier most pronounced early in neurogenesis when ribosome levels are highest, and less pronounced after mid-neurogenesis when ribosome levels decline.

Varying ribosome dwell time on a codon might be attributable to the availability of a given tRNA. Dwell time is strongly correlated with tRNA abundance in yeast ^37–39^, but is less correlated in some mammalian systems ^36,40^. We measured levels of 151 tRNA isodecoders by quantitative PCR (qPCR) array at each stage (**Extended Data Fig. 6, Supplementary Table 4**) to determine if tRNA abundance is responsible for driving ribosome dwell time differences in the developing neocortex. Usage-corrected tRNA abundance (availability) ^36^ and codon optimality – the non-uniform decoding rate between synonymous codons ^41^ – failed to show any correlation with ribosome dwell time at the A-site (**Extended Data Fig. 5c-d**).

**Fig. 6.**
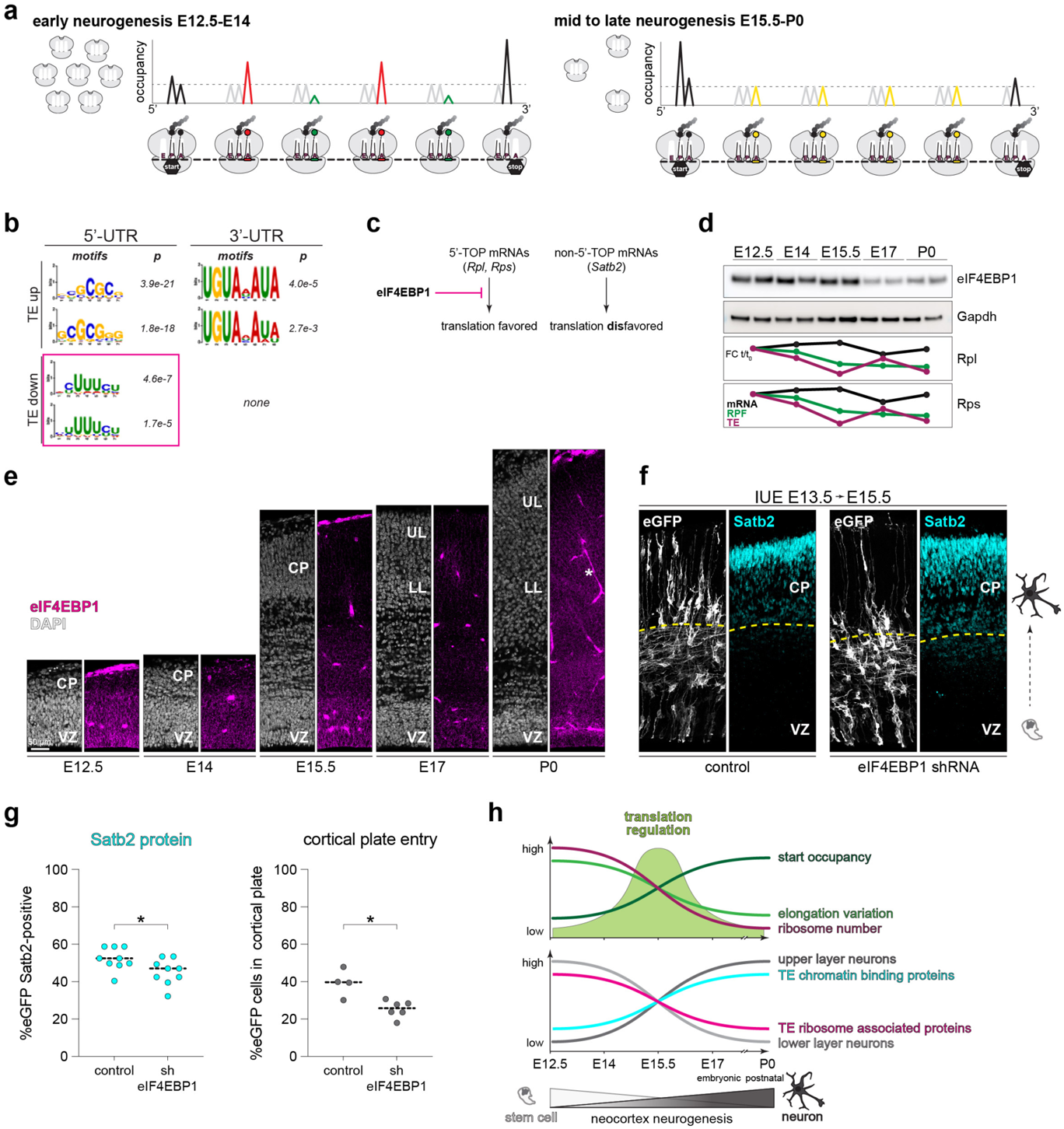
eIF4EBP1 levels coincide with ribosome abundance and control neuronal Satb2 fate *in vivo*. **a**, Model of early vs. late neurogenesis ribosome levels and per-codon changes in ribosome occupancy. **b**, Positional weight matrix of the top two motifs ranked by *p*-value in the 5’ and 3’-UTRs of TE up or down mRNAs. 5’ terminal oligopyrimidine (TOP) motifs are highlighted for TE down genes. **c**, eIF4EBP1 inhibition of ribosomal protein coding mRNA 5’-TOP sequence translation. **d**, Western blot analysis of eIF4EBP1 levels in neocortex lysates in biological duplicate (*n* = 4-6 neocortex hemispheres per lane). Concurrent trajectory of Rpl and Rps translation shown below. **e**, Immunohistochemistry analysis of eIF4EBP1 expression in neocortex coronal sections across neurogenesis. Blood vessels (star) are a common staining artifact. **f**, shRNA knockdown of eIF4EBP1 compared to scrambled control by *in utero* electroporation (IUE) at E13.5 followed by analysis at E15.5 with Satb2 protein immunolabeling. Co-electroporation of eGFP labels all transfected cells. Cortical plate neuron boundary is demarcated (dotted line). **g**, Quantification of (f) per animal for the percent of electroporated cells expressing Satb2 protein (left), and number of cells migrating into the cortical plate (right). Median (line), Mann-Whitney test, \**p* < 0.05. **h**, Summary of timed translation changes and neuronal specification during neocortex development. See also **Supplementary Table 3**.

However, we found that the amino acid coded for is a strong determinant of ribosome density occupying A-site codons, with synonymous codons showing similar occupancy (**Fig. 5g, Supplementary Table 4**). Codons for acidic amino acids are among those with the highest occupancy, suggesting they represent a kinetic barrier in early development translation ^38,42^. E12.5-E14 accounts for the extremes of A-site differences between amino acids and among synonymous codons, with a progressive, chronologic trend towards equalized occupancy by P0. Notably, some amino acids like leucine and isoleucine are coded for by both “fast” and “slow” synonymous codons, particularly apparent early in development, such as the fast TTA-Leu and slow CTG-Leu. Neither codon optimality (**Extended Data Fig. 5d**) nor codon rarity would account for such dwell time differences, as TTA-Leu is a relatively rare codon ^43^ with a short dwell time, while CTG-Leu is more common with a long dwell time.

Taken together, “fast” and “slow” amino acids in the ribosome A-site characterize the early neurogenesis period when ribosome levels are transiently abundant, while ribosome accumulation at the start codon occurs late in neurogenesis when ribosome levels decline (**Fig. 6a**). These data strongly indicate that the kinetics of translation shift sharply at mid-neurogenesis during a steep decline in ribosome levels, which coincide with major transitions in neuronal fate.

### The ribosomal protein translation inhibitor eIF4EBP1 impacts neuronal fate and migration

The overwhelming influence that changes in ribosome number can have on global protein synthesis kinetics and mRNA-specific translation is strongly supported by theoretical and experimental data ^32,33^. However, whether changes in ribosome number (**Fig. 6a**) impact neurogenesis is unknown. We first analyzed mRNAs for sequence motifs in their untranslated regions (UTRs), which are powerful regulators of neocortical translation by RNA-binding proteins ^2,14,44^. Distinct motifs are enriched in the 5’- and 3’-UTRs of mRNAs with increasing or decreasing translation efficiency (**Fig. 6b**). Translation downregulation motifs were only detected in 5’-UTRs and are enriched for terminal oligopyrimidine (5’-TOP) sequences. In translation upregulated mRNAs by contrast, 5’ GC-rich sequences and/or 3’ Pumilio binding motifs are prevalent. 5’-TOP sequences are a particular feature of ribosomal proteins coding mRNAs, and lead to their concerted translation downregulation when directly bound by their major upstream regulator eIF4EBP1 (**Fig. 6c**) ^20,21^. Since we found that ribosome levels are controlled by a timed decrease in ribosomal protein translation, we next focused on how eIF4EBP1 expression coincides with translation regulation during neocortex development.

Western blot analysis demonstrated eIF4EBP1 levels change during neocortex development, with high expression at early stages until E15.5, followed by a sharp decrease at E17, and moderate recovery at P0 (**Fig. 6d**). eIF4EBP1 levels coincide with the translation downregulation of ribosomal protein expression measured until E15.5, after which translation inhibition is transiently released at E17 (**Figs. 4b and 6d, bottom**). We next assessed the developmental expression of eIF4EBP1 *in situ* by immunohistochemistry analysis in the neocortex (**Fig. 6e**). Robust eIF4EBP1 expression was observed in neural stem cells at E12.5-E15.5 in the ventricular zone, with lower expression in cortical plate neurons. At E17, eIF4EBP1 levels decrease throughout both stem cell and neuronal zones, with some recovery at P0. These data suggest eIF4EBP1 may play a role in the early-mid neurogenic period, during a downregulation of ribosomal protein translation, which coincides with an increase in Satb2 translation.

To measure the impact of eIF4EBP1 on neuronal fate during neocortex neurogenesis, we performed shRNA knockdown by *in utero* electroporation in ventricular zone progenitors at E13.5 when eIF4EBP1 levels are high, followed by immunohistochemistry assessment of Satb2 protein expression at E15.5 when translation regulation dynamics peak (**Fig. 6f**). eIF4EBP1 knockdown in early progenitors leads to a decrease in the fraction of Satb2 protein expressing neurons at E15.5 compared to scrambled control, and arrests neuronal entry into the cortical plate (**Fig. 6f-g, Supplementary Table 3**). These data indicate that eIF4EBP1 impacts neuronal fate and migration during a large-scale transient shift in translation activity at mid-gestation.

Taken together, our data supports a model where E15.5 is a major inflection point in translation regulation during neocortex neurogenesis (**Fig. 6h**). This critical window includes a robust decrease in ribosome number in differentiating neurons, a change in translation kinetics, and global shifts in the translation efficiency of mRNAs – including genes driving neuronal specification.

### Modeling the translatome of neocortex neurogenesis

Having interrogated members of the most translationally upregulated and downregulated gene pathways, we pursued a more comprehensive bioinformatic analysis of the transcriptome-to-proteome transition in coordinated developmental programs – where deviations between mRNA and protein may represent dynamic cellular transitions ^1^. We performed hierarchical clustering of mRNA and protein expression trajectories after E12.5 per gene, which divided the proteome into 13 broad clusters (**Fig. 7a and Extended Data Fig. 7a, Supplementary Table 5**). We found clusters representing concordant and divergent trajectories between mRNA and protein, with E15.5 a common inflection point of divergent regulation. While genes with changing translation efficiency are found in all clusters, they are enriched in clusters that demonstrate highly divergent mRNA and protein expression. Furthermore, several essential neural stem cell and differentiation markers segregate into distinct clusters, such as *Nes* in cluster J and *Satb2* in cluster M. Reinforcing the biological significance of clusters representing different mRNA and protein trajectories, gene ontology analysis demonstrated many non-overlapping, distinct pathways enriched in different clusters, such as neuron differentiation processes enriched in cluster D (**Fig. 7b, Supplementary Table 5**).

**Fig. 7.**
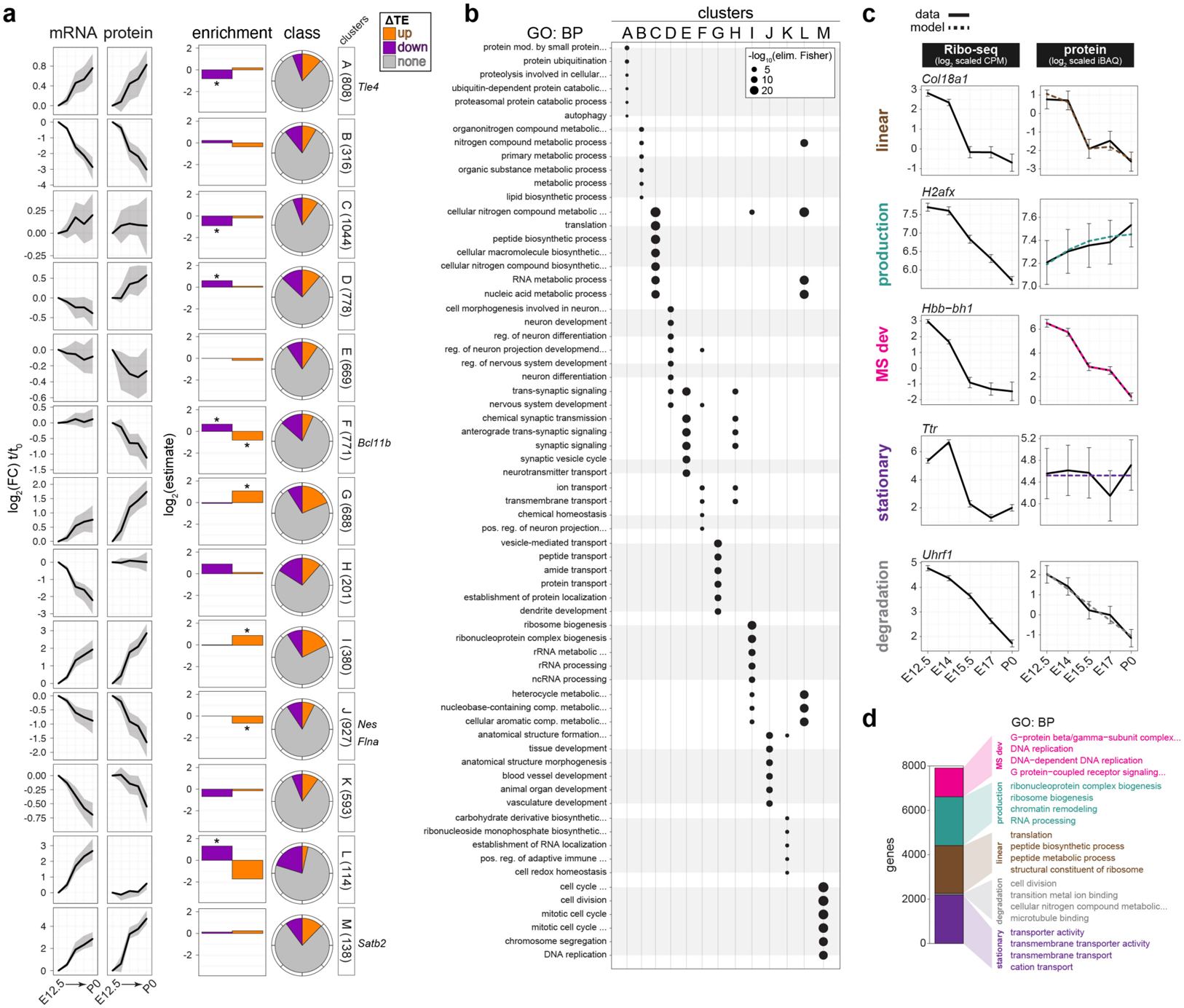
Modeling divergent trajectories of mRNA and protein expression by translation regulation. **a**, mRNA (RNA-seq) and protein (MS) expression per gene from E12.5 (t_0_) to subsequent stages (t) clustered by trajectory. The median trajectory is shown, with upper and lower quartiles (grey). Enrichment and proportion of TE up and down genes in each cluster, with significant enrichment (\**p* < 0.05). Example neural stem cell and neuronal marker genes are indicated (right). **b**, Gene ontology (GO, biological process) enrichment for each cluster, with unique terms for a cluster outlined in grey. **c**, Modeling of non-linear relationships between Ribo-seq and MS comparing active translation vs. steady-state protein, with representative genes shown for each category. See text for details. **d**, Proportion of total genes in each category from (c), with enriched GO terms per category. Fisher’s exact test, *p* < 0.05. See also **Extended Data Fig. 7, Supplementary Table 5**, and https://shiny.mdc-berlin.de/cortexomics/.

The relationship between Ribo-seq density and steady state protein levels is complicated by the fact that protein half-lives are relatively long ^45^, and reflect the cumulative effects of synthesis and degradation over time. In contrast, Ribo-seq reflects synthesis at a given time point. Deviations between protein and Ribo-seq are expected whenever protein levels have not yet reached equilibrium with synthesis, making linear comparison of protein concentrations and Ribo-seq densities difficult to interpret. We therefore made use of a kinetic, time continuous model of protein translation similar to ^46^ (see **Methods**).

We classified proteins into one of five categories (**Fig. 7c**): *stationary*, where protein levels showed little change; *linear*, where protein levels were in near-equilibrium with Ribo-seq measured synthesis; *production*, consistent with a non-equilibrium protein trajectory; *degradation*, for which protein degradation alone fit the data; and *“MSdev”*, whose protein trajectories diverged from their Ribo-seq trajectory for any combination of parameters. Then, by using the approximation of single constant relating RPF density and synthesis rate, we were able to estimate half-lives for all genes, which show a strong correlation to experimentally determined degradation rates in NIH 3T3 cells ^47^ (**Extended Data Fig. 7b**). For example, our predicted MSdev category proteins are more likely to demonstrate non-exponential decay kinetics during their lifetime (**Extended Data Fig. 7c**).

Genes in the five modeled categories showed distinct gene ontology term enrichment (**Fig. 7d**), such as the linear relationship between the translation and abundance of ribosome components, or the non-linear relationship for chromatin associated proteins. Interestingly, G-protein coupled receptors and DNA replication genes are enriched in the MSdev category, suggesting their expression patterns are highly multifaceted. In contrast, transmembrane transporter protein levels are highly stable, buffering upstream transcription/translation changes. Thus, our modeling highlights how multiple layers of post-transcriptional regulation impact distinct gene families during the time course of neuronal differentiation.

## Discussion

Our study traces how functional gene expression is catalyzed in a complex developmental system, capturing the reactants, synthesis, and products of mRNA translation across the time course of neocortex neurogenesis. We find widespread deviations in the trajectory of mRNA and protein expression along with changes in translation for ~18% of the transcriptome, with a transient peak at mid-neurogenesis. We interrogate the protein families most enriched among translation up and downregulated genes. Translation upregulation particularly impacts chromatin binding proteins like Satb2, which are essential components of neurogenesis. Translation downregulation targets the translation machinery itself, with an acute decline in ribosome number at mid-neurogenesis. The transition from relative ribosome abundance to depletion is accompanied by a chronological shift in translation processivity at the start codon and A-site amino acid during peptide elongation. eIF4EBP1, the major upstream suppressor of ribosomal protein translation efficiency, is dynamically expressed during neurogenesis in tandem with ribosome abundance, and impacts Satb2 neuronal fate specification. Finally, we model the transcriptome-to-proteome transition in neocortex development, highlighting the impact of translation in a multilayered program of neurodevelopmental gene expression.

Neural stem cells and differentiated neurons harbor a pool of mRNAs inclusive of diverse neuronal fates ^4,7^. We propose that a broad transcriptome is filtered at the protein level for tightly timed, rapidly scalable, and spatially targeted gene expression to assemble highly evolved neuronal circuits. Per gene per hour, translation is faster and more scalable than transcription by orders of magnitude ^48^, and neuronal specification transitions occur in very narrow developmental windows ^2,3^. The availability of a diverse mRNA repertoire including both lower and upper layer neuronal fates like *Bcl11b* and *Satb2*, respectively, which can be rapidly and selectively amplified by translation upregulation, is essential to specify Bcl11b or Satb2 protein exclusive neurons, in addition to Bcl11b-Satb2 double-positive neurons ^6^. Our translatome data provide a unique window into how the proteome emerges from the transcriptome in neurodevelopment (https://shiny.mdc-berlin.de/cortexomics/).

The timed decrease in ribosome number per cell in the cortical plate represents a coordinated translation downregulation of ribosomal protein synthesis. Control of ribosome number has a dominant influence on global protein synthesis kinetics and mRNA-specific translation, and can lead to “ribosomopathies” in disease states ^32,33^. With the role of eIF4EBP1 early in neurogenesis, our study joins an evolving body of work on RNA-binding proteins and ribosome cofactors that modulate protein synthesis in the developing brain ^2,5, 13–16,44^. eIF4EBP1 is a master regulator of ribosome levels by suppressing ribosomal protein synthesis ^20,21^, which we find also impacts the fate and migration of a neuronal lineage prenatally. A timed mechanism to finely tune ribosome levels may impose essential control on how and when proteins are synthesized during neuronal fate decisions.

We measure a timed, progressive developmental shift in ribosome density surrounding start and stop codons. Previous studies of the “5’ ramp” present in Ribo-seq experiments have proposed that it represents ‘slow’ synonymous codon choice near the coding sequence start – an adaptation to prevent ribosome collision further into the open reading frame ^49^. Our data argue against this as the sole mechanism of 5’ ramping, since numerous genes show an increase in start density despite the generally decreasing effect of codon choice. The increasing relative density at the 5’ of many mRNA coding sequences resembles what might be expected during a shift from ribosome abundant elongation-limited to ribosome scarce initiation-limited translation ^50^, when kinetic barriers to start codon initiation and elongation of early N-terminal peptides ^51^ become comparatively prominent. Of note, we do not observe increasing start codon density only for high translation efficiency genes, or correlation with neurite-localized translation (**Extended Data Fig. 8**). We therefore favor the hypothesis that ribosome occupancy at beginning of open reading frames becomes progressively rate-limiting for codon-independent reasons, such as scarcity of translation cofactors and ribosomal subunits later in development.

We also find that the A-site amino acid strongly influences translation speed during early neurogenesis in particular, suggesting that factors like the electrostatics of peptidyl chain elongation ^38,42^, amino acid availability, and/or tRNA aminoacylation might play a more important role in early brain development. To our knowledge, our study is the first to demonstrate differences in the fundamental nature of codon-specific ribosome density over developmental time. Our study agrees with previous work that suggest tRNA levels are not a limiting factor for translation elongation in mammals ^36,40,52^, as they are in exponentially dividing yeast ^37–39^. Notably, however, these findings do not rule out individual cases where a tRNA may influence ribosome stalling, as reported for one nervous system-specific tRNA postnatally ^53^. We measure the total tRNA pool with a protocol that does not address tRNA charging, which is a limitation of our study and an interesting future direction for investigation.

The main limitation of our study is that parallel time course measurements of the transcriptome, tRNA pool, translatome, and proteome occur in brain tissue of mixed cell types rather than single cells. In addition to scRNA-seq, tremendous advances in analysis of the single-cell translatome by Ribo-seq ^54^ were just published. While development of single-cell proteomics is still underway ^55,56^, the input requirements for tRNA measurement remain a major obstacle. At the expense of cellular resolution, we opted to perform a comprehensive analysis that enables modeling of mRNA translation in developing brain tissue. Notably, while our study is well designed to measure changes in protein synthesis, we do not measure protein degradation directly. The unexpectedly large number of “MSdev” proteins identified in our model indicates that post-translational mechanisms like degradation ^57–59^ may also have a major impact by decoupling protein and Ribo-seq trajectories, highlighting the complexity of gene regulation in the neocortex. Despite these limitations, our approach detected two important phenomena validated at the cell type-specific level – mRNA-protein uncoupling of Satb2 by translation upregulation, and coordinated translation downregulation of ribosome abundance. Bulk tissue measurements can be very informative in tandem with single cell data deconvolution in the brain ^60–62^, and we anticipate our data will be leveraged as more single cell technologies emerge.

Taken together, our data suggests a model of developmental gene expression where the levels and kinetics of translation shift during a key window of neurogenesis in the brain – creating a major inflection point of translation at mid-gestation. These developmental windows correspond to timed changes in neuronal specification from neural stem cells, where broad transcription of neuronal subtype-specific programs is ultimately refined by translational control, more precisely demarcating the boundaries of neuronal circuits in the brain.

## Acknowledgements

We thank Nadja Klein for advice regarding modeling and statistics, and Heike Heilmann for support with immuno-electron microscopy. M.L.K. would like to thank Martin Vingron for support. M.L.K. was funded by an EMBO Long-Term Postdoctoral Fellowship (190-2016), Alexander von Humboldt Foundation Postdoctoral Fellowship, and International Guest Fellowship from the Max Planck Institute for Molecular Genetics. D.H. was supported by a grant from the Klaus Tschira Boost Fund. *In utero* electroporation experiments were funded by a Russian Science Foundation Basic Research grant (19-34-51009) to V.T.

## Author Contributions

M.L.K. designed and initiated the study, with D.H. and M.C.A. making essential contributions. U.O. and V.T. supervised the study, with contributions from C.M.T.S., M.L., M.S., and T.M. Computational experiments were performed by D.H., and laboratory experiments by M.C.A., M.L.K., A.R., E.B., and R.D. Ribo-seq and RNA-seq sample preparation and sequencing were performed by U.Z. and M.L.K., and mass spectrometry sample preparation and measurement by K.I. and M.L.K. Immuno-electron microscopy samples were prepared by A.M.-W. and imaged by B.F. and M.L.K. Data were interpreted by D.H., M.C.A., and M.L.K. Manuscript figures and text were composed by D.H., M.C.A., and M.L.K., with valuable editing and input from all authors.

## Methods

### Mice

Mouse (*Mus musculus*) lines were maintained in the animal facilities of the Charité University Hospital and Lobachevsky State University. All experiments were performed in compliance with the guidelines for the welfare of experimental animals approved by the State Office for Health and Social Affairs, Council in Berlin, Landesamt für Gesundheit und Soziales (LaGeSo), permissions T0267/15, G0079/11, G206/16, and G54/19, and by the Ethical Committee of the Lobachevsky State University of Nizhny Novgorod. Mice were utilized in the embryonic (E12.5-E17) and early post-natal (P0) period, inclusive of both male and female sexes in each litter without distinction. Timed pregnant wild-type (WT) CD-1 mice utilized for Ribo-seq, RNA-seq, tRNA qPCR array, mass spectrometry, and immuno-electron microscopy were obtained from the Charles River Company (Protocol: T0267/15). Experiments with fluorescent *in situ* hybridization and immunohistochemistry were performed in NMRI WT mice. For experiments with the tdTomato reporter, *Satb2*^Cre/+^ males ^19^ were mated to NMRI wild type females (Protocols: G0079/11, G54/19, and G206/16). *Satb2*^Cre/+^ mouse genotyping was performed as described ^19^.

### Neocortex sample preparation for bioinformatics analysis

Dissection, cryogenic lysis, and determination of optical density units (ODU) were performed as described ^15^.

### Ribo-seq and RNA-seq sample preparation and sequencing

Each replicate for paired neocortex Ribo-seq and RNA-seq included 40 brains (80 hemispheres) at E12.5, 30 brains (60 hemispheres) at E14, 21 brains (42 hemispheres) at E15.5, 20 brains (40 hemispheres) at E17, and 17 brains (34 hemispheres) at P0 – performed in biological duplicate at each stage. Neocortex tissue was lysed on ice in 20 mM HEPES, 100 mM KCl, 7.5 mM MgCl_2_, pH 7.4, supplemented with 20 mM Dithiothreitol (DTT), 0.04 mM Spermine, 0.5 mM Spermidine, 1x Protease Inhibitor cOmplete EDTA-free (Roche, 05056489001), 0.3% v/v IGEPAL CA-630 detergent (Sigma, I8896) and clarified by centrifugation at 16100 x*g* for 5 min at 4 **°**C with a benchtop centrifuge. Samples were then measured for A260 ODU on a NanoDrop 1000 Spectrophotometer. Two thirds of the sample were transferred to a new tube for Ribo-seq preparation, with the remaining one third for RNA-seq was mixed with 100 U SUPERase-In RNAse inhibitor (ThermoFisher, AM2694) and frozen at −80 °C for downstream RNA isolation.

For digestion of ribosome protected RNA fragments (RPFs), Ribo-seq samples were then mixed with 60U RNAse-T1 plus 96 ng RNAse-A per ODU, and incubated for 30 min at 25 °C, shaking at 400 rpm. To stop RNAse activity, 200 U of SUPERase-In RNAse inhibitor was then added. 10-50 % 5 mL sucrose density gradients were prepared in Beckman Coulter Ultra-Clear Tubes (344057). Base buffer consisted of 20 mM HEPES, 100 mM KCl, 10 mM MgCl_2_, 20 mM Dithiothreitol (DTT), 0.04 mM Spermine, 0.5 mM Spermidine, 1x Protease Inhibitor cOmplete EDTA-free (Roche, 05056489001), 20 U/mL SUPERase-In RNAse inhibitor (ThermoFisher, AM2694), pH 7.4, prepared with 10 & 50 % sucrose w/v. Overlaid 10 & 50 % sucrose-buffer solutions were mixed to linearized gradients with a BioComp Gradient Master 107ip.

Digested lysates were overlaid on gradients pre-cooled to 4 **°**C. Gradients were centrifuged in a SW55 rotor (Beckman Coulter) for 1 hr, 4 **°**C, 37000 rpm, and fractionated using a BioComp Piston Gradient Fractionator and Pharmacia LKB SuperFrac, with real-time A260 measurement by an LKB 22238 Uvicord SII UV detector recorded using an ADC-16 PicoLogger and associated PicoLogger software. Fractions corresponding digested 80S monosomes were pooled and stored at −80 °C.

RNA isolation with TRIzol LS was then performed for both RNA-seq and Ribo-seq samples, as per the manufacturer’s instructions. For Ribo-seq and RNA-seq samples, downstream library preparation and sequencing were performed as described ^15^. RNA-seq data were utilized in a recent study ^15^ corresponding to NIH GEO entry GSE157425. Ribo-seq data in this study are deposited in the NIH GEO: GSE169457.

### Mass spectrometry sample preparation

Total proteome analysis from neocortex lysates at E12.5, E14, E15.5, E17, and P0, including complete lysis in RIPA buffer, and downstream processing for mass spectrometry analysis, was performed in a recent study ^15^ corresponding to ProteomeXchange entry PXD014841.

### tRNA qPCR array sample preparation and measurement

tRNA qPCR array measurement of 151 tRNA isodecoders was performed by Arraystar (Maryland, USA) for neocortex lysates at E12.5, E14, E15.5, E17, and P0 from the same total RNA isolated for RNA-seq described above (**Supplementary Table 4**). Data are deposited in the NIH GEO: GSE169621.

### Ribo-seq and RNA-seq data processing

Raw sequence data was converted to FASTQ format using bcl2fastq (https://support.illumina.com/downloads/bcl2fastq-conversion-software-v2-20.html). Adapters (sequence TGGAATTCTCGGGTGCCAAGG) were removed from Ribo-seq reads with cutadapt ^63^, as well as sequences with a quality score less than 20 or a remaining sequence length less than 12, and after removing duplicate read sequences, 4bp UMIs were trimmed from either end of each sequence using a custom perl script. Ribo-seq reads were then aligned to an index of common contaminants (including tRNA, rRNA, and snoRNA sequences) using bowtie2 ^64^. The resulting processed read files were then aligned to coding sequences (the pc_transcripts fasta file), and separately, to the genome, from GENCODE release M12 (*Mus musculus*) using STAR ^65^, with the following settings: STAR --outSAMmode NoQS -- outSAMattributes NH NM --seedSearchLmax 10 --outFilterMultimapScoreRange 0 -- outFilterMultimapNmax 255 --outFilterMismatchNmax 1 --outFilterIntronMotifs RemoveNoncanonical. RNA-seq and Ribo-seq libraries achieved high coverage, with a median of 33M and 12M reads mapped to protein coding transcripts, respectively. For quality control, downstream analysis focused on coding sequences with 32 or more Ribo-seq footprints in at least one stage as per ^24^, which resulted in a set of 12,228 translated gencode transcripts (**Supplementary Table 1**).

Linear fold changes for RNA-seq and Ribo-seq were calculated using limma ^66^, for TE was calculated using xtail ^67^, and for MS calculated using proDA (https://github.com/const-ae/proDA) (**Supplementary Table 2**).

Since ribosomes with their A-site over a given position will produce a distribution of read lengths mapping to nearby positions, A/P-site alignment represents a crucial step in the processing of Ribo-seq datasets. Frequently, algorithms for A-site alignment rely either explicitly ^37,68^ or implicitly (Ahmed et al., 2019;) on the presence of large peaks at the start and/or stop codons, the known location of which provides a ‘true positive’ that can be used to choose P-site offsets for each read length. We found that such methods gave inconsistent results in our data, with optimal P-sites being chosen at biochemically implausible values (e.g., at 0 base pairs from the read 5’-end). This is likely due to 1) the variable occupancy of the start/stop peak in our data, and 2) the presence of cut-site bias in our data due to the necessity of RNAse T1 & A digestion. Calculating RUST scores and ‘metacodon’ ^35^ plots of RPF 5’-end occurrence showed that the most variation between different codons and time points (other than cut-site bias itself at RPF termini) was nonetheless limited to a narrow region a consistent distance from the codon, for each read length. Plotting KL-Divergence between observed and expected RUST scores at different distances from the read 5’ end and measuring the between-codon variance at each position revealed that it aligned with an offset of approximately 14-15 nt (consistent with the A-site position) for reads of length between 25 and 31, and so we chose these for further analysis of ribosome dwell time (**Supplementary Table 4**). We also observed an adjacent region of lesser variability 3bp towards the RPF 5’ end, consistent with a non-zero but significantly less influence of the P-site codon (**Supplementary Table 4**).

The program DeepShapePrime ^70^, modified to accept our chosen P-site offsets instead of hardcoded ones, was then used to derive isoform specific abundance measurements for each protein coding transcript.

In parallel to the above, iso-form level quantification of the RNA-seq was carried out using salmon ^71^, with an index built from coding M12 sequences, and the following settings: salmon quant -l SR --seqBias --validateMappings

A snakemake ^72^ file automating the above workflow is available at: https://github.com/ohlerlab/cortexomics.

We then converted DeepShape-prime’s output to salmon format to combine both outputs, using the ORF length as effective length. The R package tximport ^73^ was used to derive length-corrected gene-level counts and isoform level counts and TPMs for both datasets. The voom package was used for variance stabilization and linear modelling of this data to derive confidence intervals for transcriptional and translational change, both relative to E12.5, and stepwise between each stage. The xtail package ^67^, which is specifically geared towards estimation of translational efficiency (TE; i.e. the ratio of Ribo-seq density to RNA-seq density) change in the presence of transcriptional change, was used to detect changing TE. Numbers for TE change quoted in the text refer to xtail’s differential TE calls, with stepwise fold changes shown in **Fig. 1**, and TE changing genes being elsewhere defined relative to E12.5 (**Supplementary Table 2**).

For metagene plots, a ‘best transcript’ (the transcript with the highest median Ribo-seq coverage across all samples) was selected for each gene. These transcripts were further limited to those with a length of 192 or greater. Each of these transcripts was also also analyzed using the RiboDiPA ^34^ package, which looks for position-specific differences in Ribo-seq occupancy between conditions. Since metacodon plots indicated that changes at the start and stop codon were limited to a distinct region 3-4 codons from the start and stop, we divided each coding sequence in to 15 bins, with 7 bins of 4 codons each centering on the start and stop, and a final ‘mid’ bin of variable size encompassing the rest of the ORF (ORFs too short to accommodate this were excluded). We then plotted bin-level log2 fold changes for each gene with significant q-value of using the AUG/stop changing bins.

Fold changes were binarized into ‘significant’ (absolute fold change greater or less than 1.25, adjusted p-value < 0.05) and ‘non-significant’ for plotting up and down regulated genes, respectively, and GO term analysis – referred to as dTE and non-dTE in the case of TE fold-change. For GO term analyses of TE change and positional Ribo-seq change, The R package topGO was used.

A list of ribosomal proteins for the mouse large and small subunits were curated from Uniprot.

### tRNA abundance and codon dwell time analysis

tRNA abundance was calculated from Arraystar™ Ct values by the negative delta Ct value for each tRNA compared to the mean of 5S and 18S rRNA levels in each sample (**Supplementary Table 4**). Abundance per-codon was calculated by taking the mean of each replicate, and summing values for all relevant iso-decoders. Availability ^36^ was calculated as the residual from a simple linear model regressing codon usage against abundance, where codon usage was defined as the occurrence of that codon in the M12 coding transcriptome, weighted by the relevant TPM of each transcript in that sample. We attempted weighting by wobble base pairs as in ^41^ and found this did not impact the conclusions.

We followed the approach of ^35^ and used RUST values as a robust estimator of codon specific dwell times. A-site occupancy was defined as the RUST value for that codon at the point of maximum variation (14 or 15 base pairs from the 5’ end) with P-site occupancy defined as the RUST value 3 bp closer to the 5’ from there (**Supplementary Table 4**)

Relationships between codon dwell time, tRNA abundance/availability, and amino acid identity, were investigated using the R function lm. The dataset used consisted of 269 – (i.e. one per quantified codon, per sample) with terms for the stage of the sample (S), the amino acid coded for (AA), and the abundance (or availability) of the encoding tRNAs (AB).

The largest explanatory variable was AA, which also showed a significant interaction with S, indicating that the amino acid coded for explained ~ 34% of the variance in dwell time between codons. This term also showed a significant interaction with sample stage, indicating that the amino-acid specific factors determining dwell time may vary over development (e.g., due to the availability of amino acids changing). Within a sample or across all samples, there was no association between tRNA abundance and dwell time, even after correcting for the effect of amino acid coded-for. Some codons however show a significant interaction between abundance and developmental stage, and because these codons were biased towards the high or low end of the abundance dwell time spectrum, we plotted time-relative change vs. abundance, for the top and bottom quartiles of dwell time abundance. This revealed a significant association between change in time-relative tRNA abundance and dwell time, with fastest codons showing decreasing tRNA abundance as they slowed, and the slowest codons also showing decreasing tRNA abundance.

### Mass Spectrometry data processing

All raw data were analyzed and processed by MaxQuant (v1.6.0.1) ^74^ (**Supplementary Table 1**). Default settings were kept except that ‘match between runs’ was turned on. Search parameters included two missed cleavage sites, cysteine carbamidomethyl fixed modification and variable modifications including methionine oxidation, protein N-terminal acetylation and deamidation of glutamine and asparagine. The peptide mass tolerance was 6ppm and the MS/MS tolerance was 20ppm. Minimal peptide length of 7 amino acids was required. Database search was performed with Andromeda ^75^ against the UniProt/SwissProt mouse database (downloaded 01/2019) with common serum contaminants and enzyme sequences. The false discovery rate (FDR) was set to 1% at peptide spectrum match (PSM) level and at protein level. Protein quantification across samples was performed using the label-free quantification (LFQ) algorithm ^76^. A minimum peptide count required for LFQ protein quantification was set to two. Only proteins quantified in at least two out of the three biological replicates were considered for further analyses.

To improve the match between mass spec data and sequence data, the peptides from each mass spec group were matched against M12 protein sequences. Instances in which a UNIPROT gene identifier did not match any gene in GENCODE, but in which the associated peptide sequences matched proteins for a single GENCODE gene, were updated to match that Gencode gene. All further analyses were carried out using gene-level proteomic data.

The R package proDA was used to calculate dropout-aware abundance estimates for each protein group, as well as fold changes and confidence intervals relative to E12.5. For each gene, a ‘best’ matching protein group was defined as the one with the least missing, and highest median, signal across all samples, and selected for further analysis.

### Analysis of variance

Analysis of variance was carried out a manner similar to ^25^. We fit a linear model regressing measured protein levels, or protein fold changes, P, against measured Ribo-seq or RNA-seq levels R. We then performed variance decomposition using the following equation:

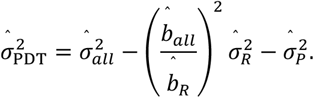

Where 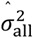 represents total variance in measured protein abundance, (i.e. in proDA-normalized LFQ values) and is decomposed into stochastic error in protein measurement 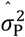 (estimated standard error of the protein abundance model fit using proDA), systematic variation in protein levels independent of 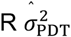, and error in R measurement, where 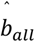 is the linear coefficient relating Ribo-seq and RNA-seq measurements to protein abundance, 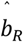 is the measurement bias for R, and 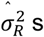 the stochastic measurement error in R. Lacking a means of measuring 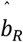 in our data, we experimented with a range of values, including the experimentally determined value of 1.21 based on NanoString measurements by ^34^. We found that due to the relatively minor stochastic error in measurements of R, our estimates of 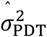 were robust to reasonable values of 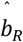 (between 0.75 and 1.5) and so we elected to fix its value at 1. We then calculated variance explained as:

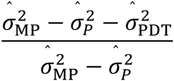

We applied this equation both within each time point, and to the fold changes between each time point. Stochastic error terms for both within-stage and between stage values for R and P were calculated using limma and proDA respectively. Notably, correlation between the two sequencing assays and MS is strongly dependent on the magnitude of change at that time point, with technical noise specific to each assay non-correlated ^1^. For the R implementation of the above equations, see our github repository (https://github.com/ohlerlab/cortexomics), and the file src/Figures/Figure4/2_vardecomp.R.

### Hierarchical clustering

For hierarchical clustering (**Supplementary Table 5**), we took fold changes in RNA-seq and MS values relative to E12.5, for each gene, and carried out PCA on the resulting n x 8-dimensional matrix. We calculated Euclidean distances between genes and performed hierarchical clustering using the R function *hclust* and the ‘ward’ clustering criterion – i.e., favoring the creation of large clusters rather than small clusters containing few outliers. We found that our expression data showed a smooth reduction in variance explained as the number of clusters varied, and so we plotted GO-term enrichment for different cluster numbers, and finding that clusters with similar GO-term enrichments began to appear at a cluster number of 13 chose this as our cutoff. Meta-trajectories for each cluster were plotted using the median and upper/lower quartiles for each cluster. Enrichment of dTE genes in each cluster was calculated using Fisher’s exact test (with dTE status, and inclusion in the cluster, as binary variables). GO term analysis of each cluster was carried out using topGO.

### Nonlinear trajectory modeling

In order to model the nonlinear relationship between steady-state protein levels and Ribo-seq, a measure of protein synthesis, we used an approach similar to that used by ^46^ -our full ‘*production’* model represents the expression of each protein as the result of a synthesis rate, directly proportional to Ribo-seq footprint density with a proportionality constant Ks (or SDR - see ^38^), and a decay rate Kd:

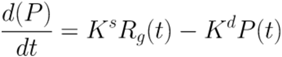

If the functional form of Ribo-seq density is modeled as a linear stepwise function, this equation has an analytic solution ^46^. In practice, the parameters Ks and Kd will be non-identifiable depending on the trajectory shape and half-life of the protein involved; for many proteins, only their ratio, defining the equilibrium steady state, can be estimated (along with the initial value of P). In addition to the ‘*production’* model, we included reduced versions of our model which fixed Kd at a high value (the ‘*linear’* model) giving a linear protein-Ribo-seq relationship, fixed Ks at a low value and modeled protein as controlled by degradation only (the *‘degradation’* model), or fixed both to leave protein levels stationary (the *‘stationary’* model). We further included a model allowing arbitrary deviations from the Ribo-seq trajectory (the *‘MSdev’* model), since many proteins showed changes in their trajectory that were not explicable by any value of Ks and Kd. We used the bayesian information criterion (BIC) to select an optimal model for each gene, further requiring that residuals in this model be normally distributed (as per a chi-squared test). To estimate half-lives, we made the simplifying assumption of a single Ks value applying to all genes, allowing pi-half estimates to be derived for all proteins.

Stan files detailing the above models are available on the project github, and data are in **Supplementary Table 5.**

### Single cell RNA-seq data

Single cell RNA-seq (scRNA-seq) data was derived from data and scripts in ^4^ and accompanying web resource: http://genebrowser.unige.ch/telagirdon/#query_the_atlas.

For each gene, it’s occurrence in neocortex cells measured by scRNA-seq is presented as a heat map arranged by chronological time of cell collection (x-axis) vs. time since cell birth (y-axis), after a timed pulse with a FlashTag label *in utero*. These axes correspond to roughly orthogonal programs of gene expression change, with the y-axis describing differences between apical progenitors and differentiated neurons, and the x-axis describing differences between cells born at different stages of development.

### Sequence motif analysis

Motif analysis was performed with the AME program from the Meme suite as per ^46,77^ because we observed a systematic difference in UTR length between TE changing and TE unchanging genes. AME requires that input and control sequences are of approximately equal length distribution, so we created a sample of TE changing genes whose length distribution matched that of the TE unchanging genes. We ran AME with the CISBP-RNA database of RNA-binding protein motifs ^78^.

### Immuno-electron microscopy

Fixation, sectioning, immunolabeling, and electron microscopy were performed as described previously ^15^. E12.5 and E15.5 neocortex coronal sections were labeled with mouse anti-Rps5 (uS7; Santa Cruz, sc-390935) followed by 2.5 nm nanogold conjugated secondary antibody (Nanoprobes, 2001). Imaging was performed at 2700 x magnification on a Tecnai Spirit electron microscope. Quantification was performed in FIJI ^79^ with the Process > Find Maxima tool, and Measure > Area tool, followed by statistical analysis in GraphPad Prism (GraphPad Software Inc.) to calculate puncta per μm^2^ (Welch’s ANOVA, Dunnett’s *post hoc* test) (**Supplementary Table 3**). Primary antibody leave-out controls were prepared in parallel, and were absent of nanogold signal.

### Expression vectors

For tdTomato reporter experiments, we used beta-actin driven expression constructs *pCAG-EGFP* and *pCAG-flox-STOP-flox-tdTomato*, as described previously ^31^. A control vector with scrambled non-silencing shRNA ^59^ was obtained from Thermo Scientific, and the shRNA to knock-down plasmid for mouse *eIF4EBP1* was obtained from Sigma Aldrich (TRCN0000335381).

### *In utero* electroporation (IUE)

Mouse embryos were subjected to IUE exactly as described previously ^31,59,80^. For the experiments with the tdTomato reporter in the Satb2^Cre/+^ line, we used an equal amount of *pCAG-GFP* and *pCAG-flox-stop-flox-tdTomato*.

### Fluorescent *in situ* hybridization (FISH)

*In situ* hybridization using RNAscope Technology to detect mRNA of *m. musculus Satb2* (413261) and *Bcl11b* (413271-C2) was performed according to the manufacturer’s protocols (ACD, 323100). Prior to hybridization, embryonic brains at E12.5, E14.5 and E16.5 were collected in PBS, fixed in 4 % PFA/PBS prepared with DEPC for 16-20 hours at 4 °C. Brains were then incubated in sucrose solutions (10 % - 20 % - 30 %/PBS) until they reach osmotic equilibrium, embedded in O. C. T. Compound (Tissue-Tek) in a plastic cryoblock mold and frozen on dry ice. Coronal sections of 16 μm thickness were collected using a cryostat.

### Cryosectioning

For all histological procedures, brain sections were prepared on a Leica CM3050S cryostat. Prior to cryosectioning, brains were incubated for at least 5 hours with 10% sucrose in PBS, followed by incubation with 30% sucrose in PBS until the tissue reached osmotic equilibrium. Next, brains were frozen in −38 to −40°C isopentane (Roth). For processing of the tissue after *in utero* electroporation, coronal cryosections of 50 μm thickness were collected in PBS/0.01% sodium azide solution. For *in situ* hybridization and the mRNA/protein colocalization experiments, 16 μm sections were collected.

### Immunohistochemistry

Fixed brain sections were washed with PBS three times at room temperature prior to the procedure to remove the sucrose and freezing compound residue. The sections were then incubated with Blocking solution (5% goat serum, 0.5% (v/v) Triton X-100, PBS) for one hour at room temperature, then with the primary antibody and DAPI diluted in blocking buffer for 16-20 hours at 4 °C, washed in PBS three times for 30 minutes and incubated with secondary antibody diluted in the blocking buffer for up to four hours at room temperature. Next, sections were incubated with PBS for 30 minutes three times and mounted with a cover glass (Menzel-Gläser) and Immu-Mount mounting medium (Shandon, Thermo-Scientific). For experiments with dual mRNA and protein labeling, instead of mounting after the hybridization protocol, the sections were subjected to the immunohistochemistry as described here.

### Antibodies for immunohistochemistry

Primary antibodies used for immunocytochemistry were used at dilutions indicated: anti-Satb2 (1:500, rabbit, home-made; ^31^), anti-Bcl11b (1:500, rat, Abcam, 25B6, anti-Ctip2, RRID:AB_2064130), anti-GFP (1:1000, goat, Rockland, RRID:AB_2612804), anti-Cre (1:1000, rabbit, SySy, RRID:AB_2619968), anti-Tbr2 (1:300, rabbit, Abcam, RRID:AB_778267), anti-Pax6 (1:500, rabbit, Millipore, RRID:AB_1587367), Draq5 (1:2000), anti-eIF4EBP1 (1:1000, rabbit, Abcam, ab32024, RRID:AB_2097990. All secondary antibodies were from Jackson Immunoresearch and were used at 1:250.

### Confocal imaging

Imaging of brain coronal cross sections after IUE was performed at the level of primary somatosensory cortex primordium. For imaging of the overview of immunostaining, a Leica SPL confocal microscope with 20X, 40X and 63X objectives was used. For quantitative imaging of FISH signal, a Leica Sp8 microscope with 40X objective was used. Quantification of mRNA cluster sizes, and mRNA and protein localization, was performed using ImageJ software.

### Quantification of distribution and size of mRNA clusters

mRNA puncta were quantified using ImageJ software. The maximum intensity of confocal image Z-stacks was projected on a single 2D plane. After thresholding, the images were binarized using the watershed segmentation to separate cluster clouds. The number of particles of 0.1 μm^2^ or bigger were then quantified using Measure Particles tool and normalized to the number of DAPI-labeled nuclei in a given cortical area (VZ, CP, etc.). Area of clusters was quantified as well and expressed as an absolute surface. See **Supplementary Table 3.**

### Quantification of colocalization

Mander’s colocalization coefficient was quantified for neurons expressing Satb2 and Bcl11b protein and mRNA. Protein colocalization was determined manually, and RNA colocalization was quantified using binarized images after multiplication. See **Supplementary Table 3.**

### Quantification of neuronal cell markers

The manually quantified number of neurons expressing a given marker was normalized to the entire number of IUE-labeled neurons or to DAPI-labeled nuclei count. See **Supplementary Table 3.**

### Statistical analyses

Statistics were performed using SPSS v.17 (San Diego, USA) or GraphPad Prism software. All numerical values and description of statistical tests used, definition of center, dispersion, precision, and definition of significance can be found in **Supplementary Table 3**. Prior to comparison of experimental groups, normality and log-normality test were performed.

### Data availability

Code generated during this study is supplied at: https://github.com/ohlerlab/cortexomics. Further requests may be directed to and will be fulfilled by the Lead Contact, (M.L.K.). Data have been deposited in publicly available repositories as indicated:

RNA-seq data are publicly available in the NIH GEO: GSE157425.

Ribo-seq data are deposited in the NIH GEO: GSE169457.

tRNA qPCR array data are deposited in the NIH GEO: GSE169621.

Mass spectrometry data are publicly available in the ProteomeXchange: PXD014841.

## Extended Data

**Extended Data Fig. 1.**
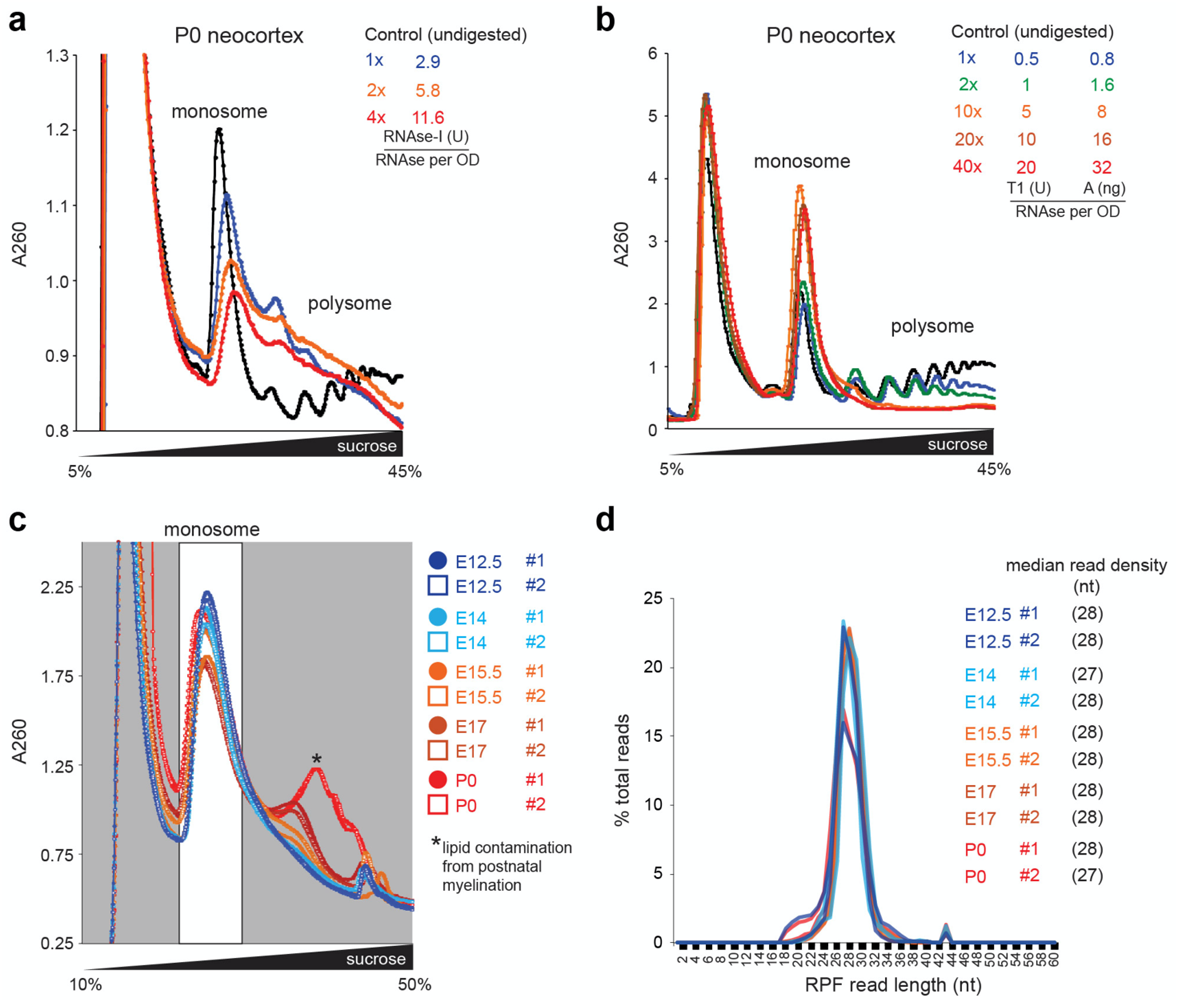
Optimized ribosome protected mRNA fragment purification from neocortex. Nuclease digestion for the generation of ribosome protected mRNA fragments (RPFs) from P0 neocortex, with **a**, RNAse-I vs. **b**, a combination of RNAse-T1 & A. Absorbance at 260 nm (A260). Chains of actively translating ribosomes (polysome) should be digested into single ribosomes (monosome). RNAse-I, typically used in yeast, was inefficient in neocortex lysates, and thus an RNAse-T1 & A protocol was used for this study. **c**, Nuclease digestion and purification of neocortex RPFs in biological duplicates at each developmental stage with the optimized protocol from (b). Each biological replicate included 17-40 brains (34-80 neocortex hemispheres) as detailed in the **Methods**. **d**, RPF read length distribution. Associated with Fig. 1. See also **Supplementary Table 1**.

**Extended Data Fig. 2.**
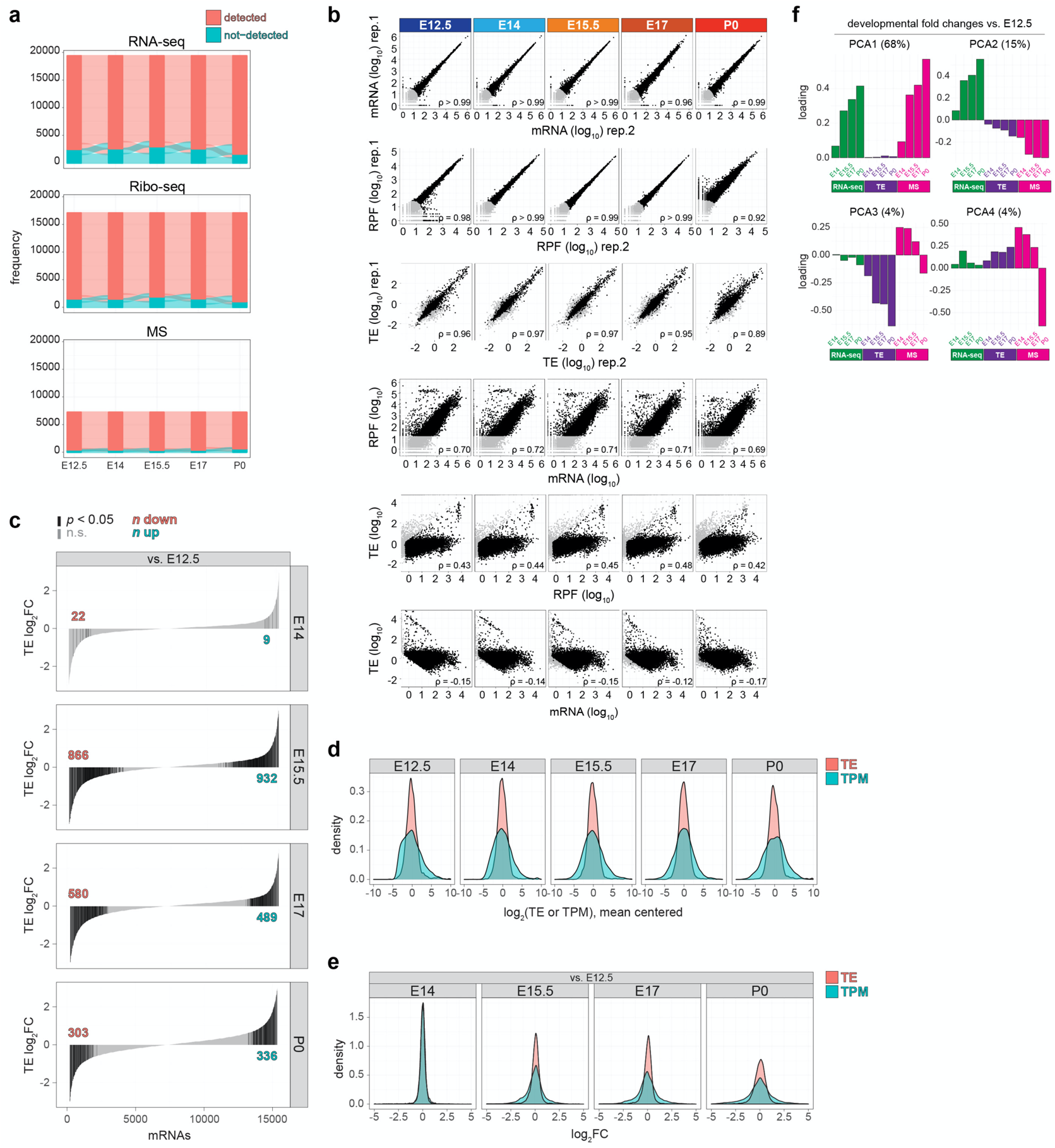
Neocortex RNA-seq, Ribo-seq, MS, and translation efficiency data characteristics. **a**, River plots demonstrating the number of unique genes detected across all 5 stages measured by RNA-seq, Ribo-seq, or mass spectrometry, compared to the number detected in <5 stages. **b**, Biological replicates of transcripts per million (TPM) measured by RNA-seq (mRNA), Ribo-seq (RPF), and calculated translation efficiency (TE), including correlations between RPF and TE with mRNA to highlight genes with robust translation regulation. **c**, The distribution of TE up and down fold changes (FC) compared to the earliest stage E12.5, with significant genes highlighted in black (*p* < 0.05). **d**, The distribution of TE and mRNA abundance (TPM) for all genes at each stage, and **e**, fold changes vs. the earliest stage E12.5. **f**, Principal component analysis (PCA) of developmental fold changes in RNA-seq, TE, and MS compared to the earliest stage E12.5. The first four components are shown, with percent variance annotated. Associated with Fig. 1. See also **Supplementary Tables 1-2**.

**Extended Data Fig. 3.**
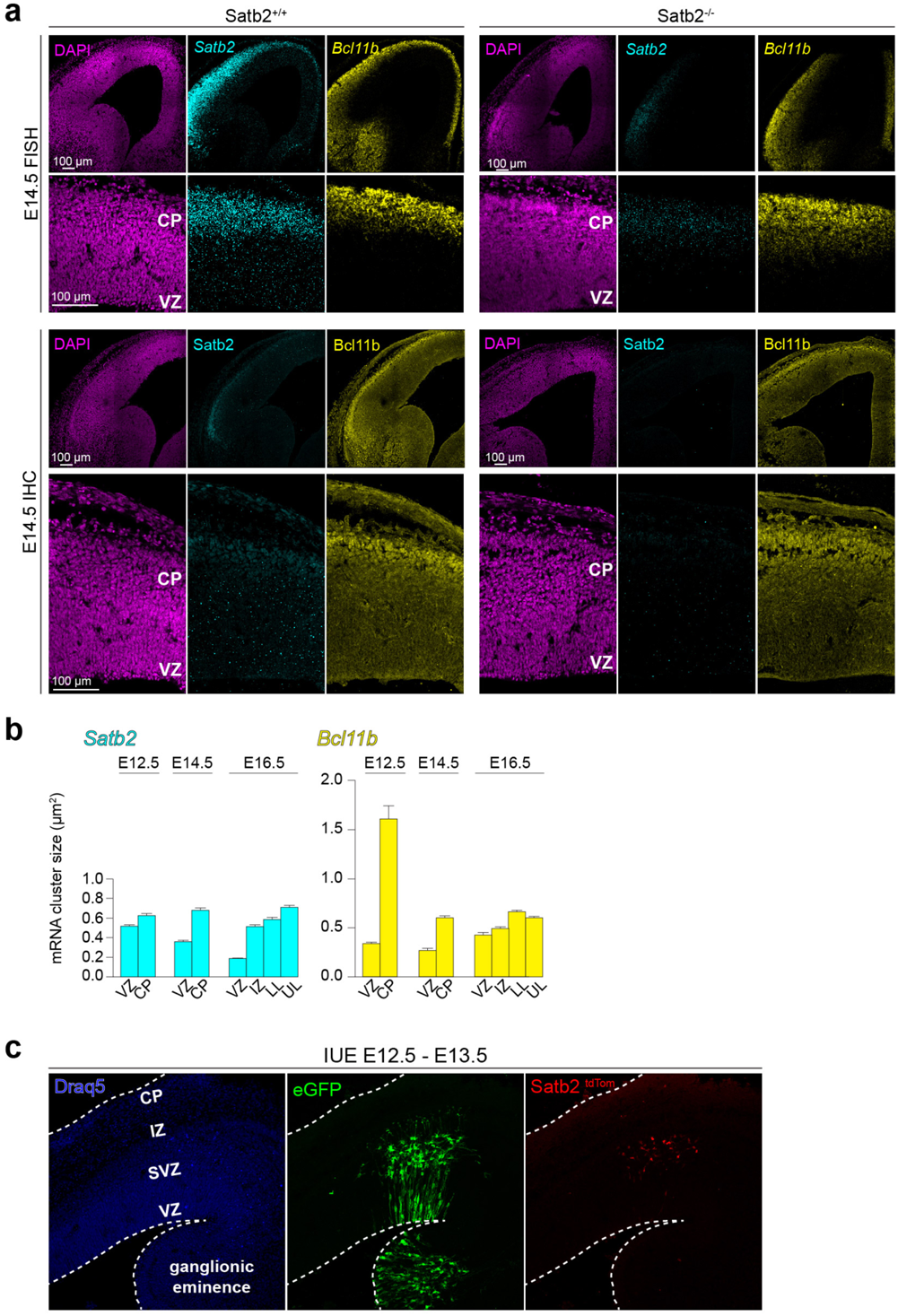
*Satb2^-/-^* control for FISH and IHC and neocortex-specific *Satb2* transcription. **a**, Fluorescence *in situ* hybridization (FISH) and immunohistochemistry (IHC) probing for *Satb2* and *Bcl11b* mRNA and protein, respectively, in wild-type (*Satb2^+/+^*) and *Satb2* knockout (*Satb2^-/-^*) neocortex coronal sections at E14.5. Ventricular zone (VZ), cortical plate (CP). **b**, Measurement of *Satb2* and *Bcl11b* mRNA cluster sizes in FISH probed neocortex sections at three developmental stages. Intermediate zone (IZ), lower layers (LL), upper layers (UL). Mean ± SEM. **c**, *Satb2* transcription activation visualized in *Satb2^Cre/+^* mice by *in utero* co-electroporation of the neocortex and ganglionic eminence with a *loxP-STOP-loxP-tdTomato* (*Satb2^tdTom^*) fluorescence reporter at E12.5, along with eGFP reporter for all transfected cells, and analysis in coronal sections at E13.5. Sub-ventricular zone (SVZ). Associated with **figED. 2-3**. See also **Supplementary Table 3**.

**Extended Data Fig. 4.**
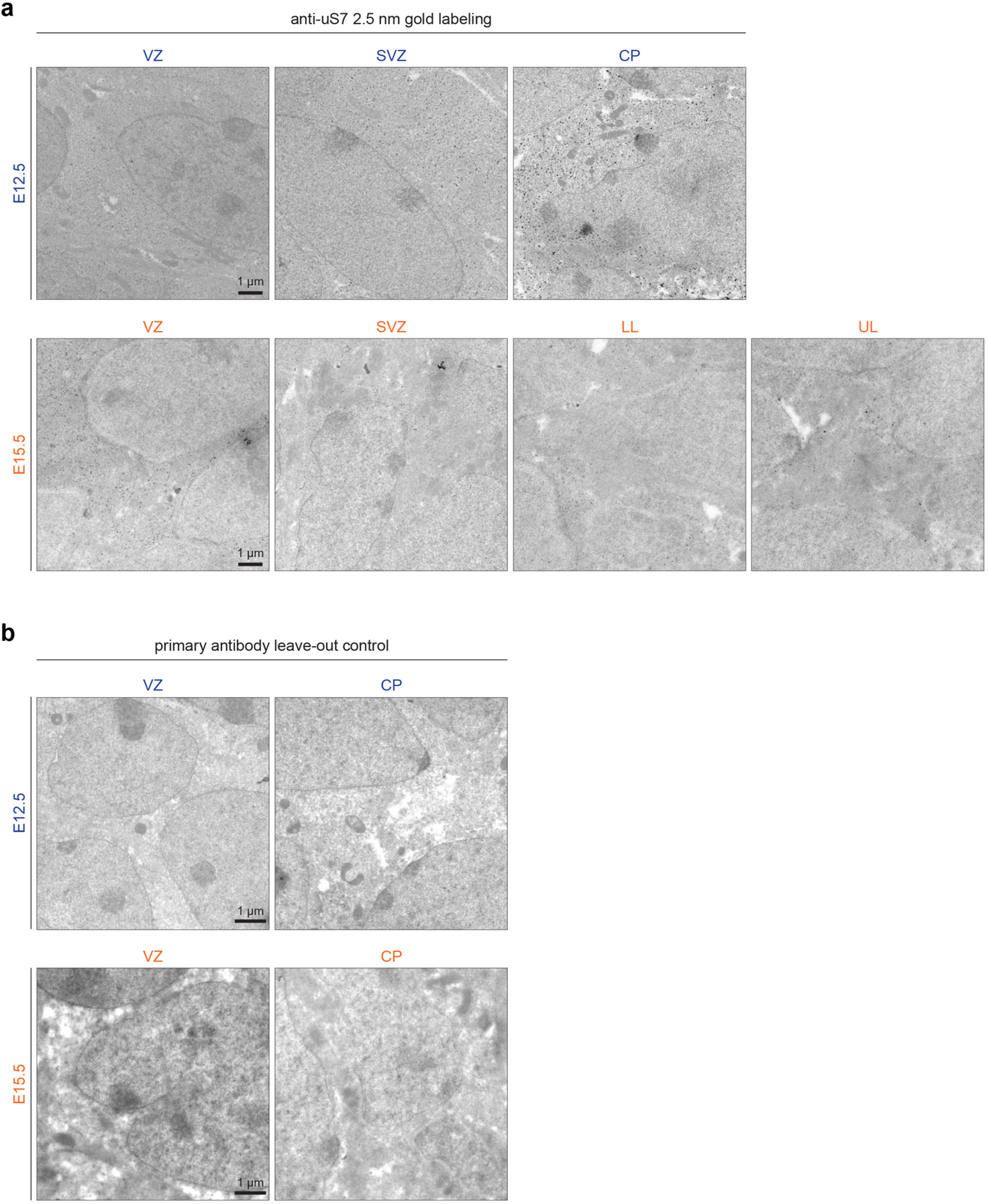
Immuno-electron microscopy labeling of ribosomes. **a**, Raw images of neocortex coronal sections at E12.5 and E15.5 shown in Fig. 4c, immunolabeled with anti-ribosomal protein uS7 followed by 2.5 nm gold secondary (dark black spots), which were automatically detected and quantified in FIJI (magenta spots in Fig. 4c). Electron microscopy was performed in regions corresponding to the stem cell niches of the ventricular zone (VZ) and sub-ventricular zone (SVZ), in addition to regions of differentiating neurons in the cortical plate (CP), which includes both lower layers (LL) and upper layers (UL) at later stages. Quantification of nanogold secondary signal was performed per unit area of the cytoplasm, with nuclei excluded by tracing the nuclear membrane (black lines in Fig. 4c). **b**, Primary antibody leave-out controls were prepared in parallel.

**Extended Data Fig. 5.**
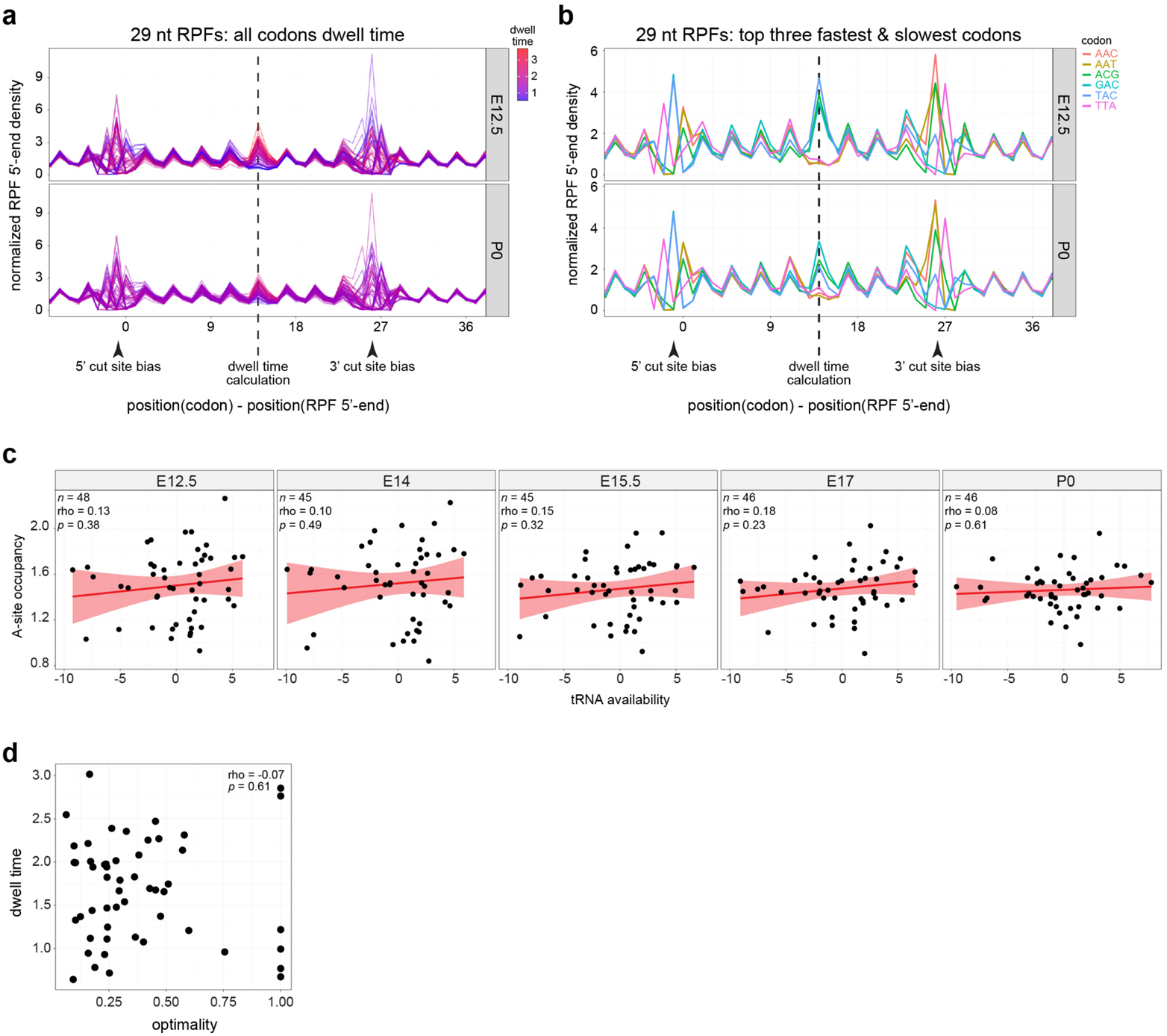
Analysis of per-codon ribosome density. 5’ normalized ribosome-protected mRNA fragment (RPF) density for **a**, all codons and **b**, the top 3 slowest/fastest codons. Plotting the normalized density of Ribo-seq read 5’ ends relative to each codon/read length/sample shows two strongly variable regions corresponding to 5’- and 3’-end cut site biases during nuclease digestion. A third variable region in between corresponds to RPFs with their A/P-sites positioned over the codon. We infer the location of the A-site as the 3 bp region showing the most inter-codon variability, and use the normalized occupancy here to measure codon dwell times, and variance between codons. Independently, this region also identifies the location of intra-codon variability between samples. **c**, Per-codon correlation between tRNA availability calculated from tRNA qPCR array (see **Methods**), and the ribosome occupancy of that codon when positioned in the A- or P-site of the ribosome footprint. **d**, Correlation between ribosome dwell time per codon and the optimality of the codon as defined in (dos Reis et al., 2004), with the mean across all stages shown. Associated with Fig. 5. See also **Supplementary Table 4**.

**Extended Data Fig. 6.**
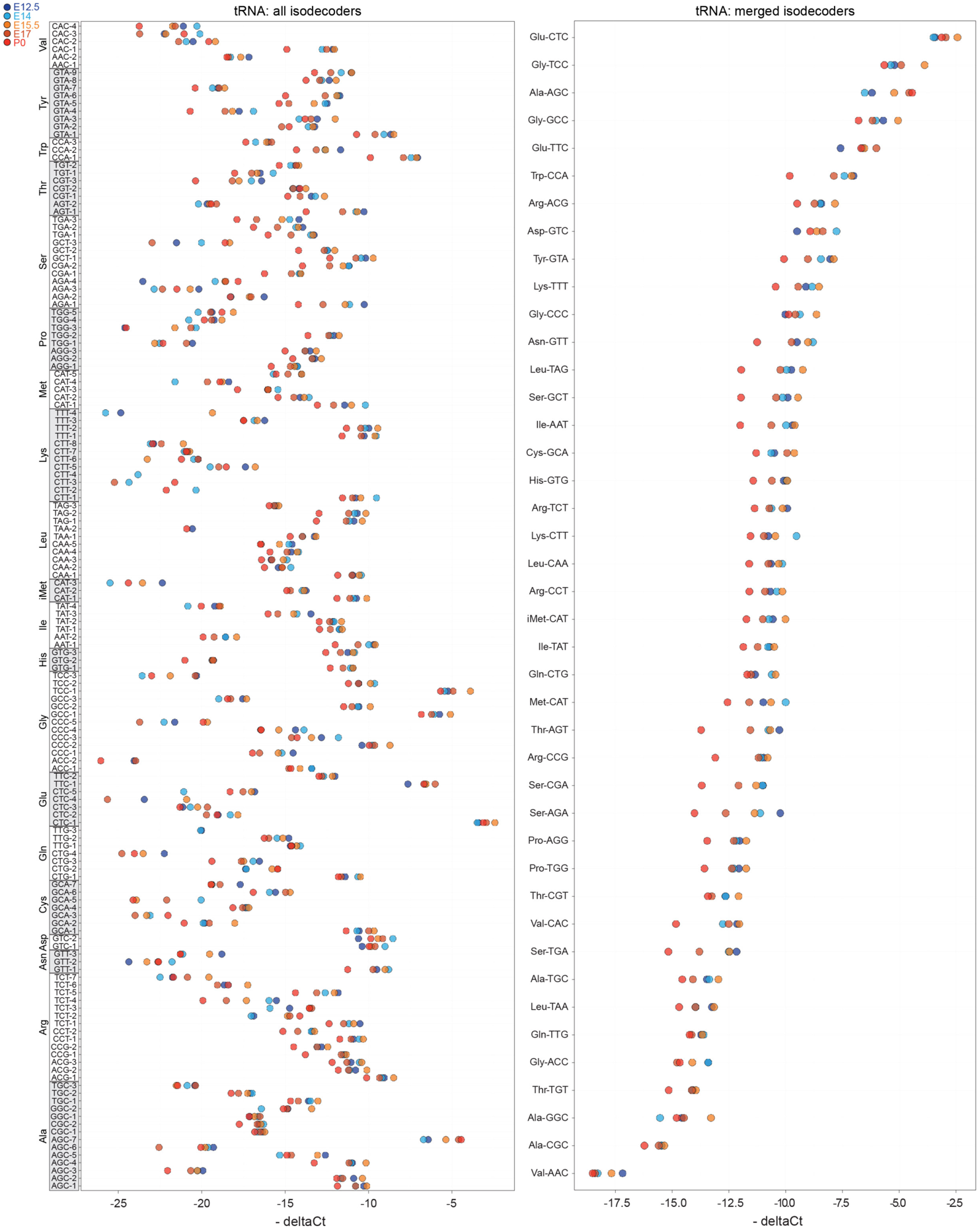
Neocortex tRNA qPCR array. Total tRNA levels at each stage measured by qPCR array in biological duplicate, with Ct values for each tRNA isodecoder (left) or averaged across isodecoders (right) compared to the mean of 5S and 18S rRNA levels in each sample (delta Ct). Associated with Fig. 5. See also **Supplementary Table 4**.

**Extended Data Fig. 7.**
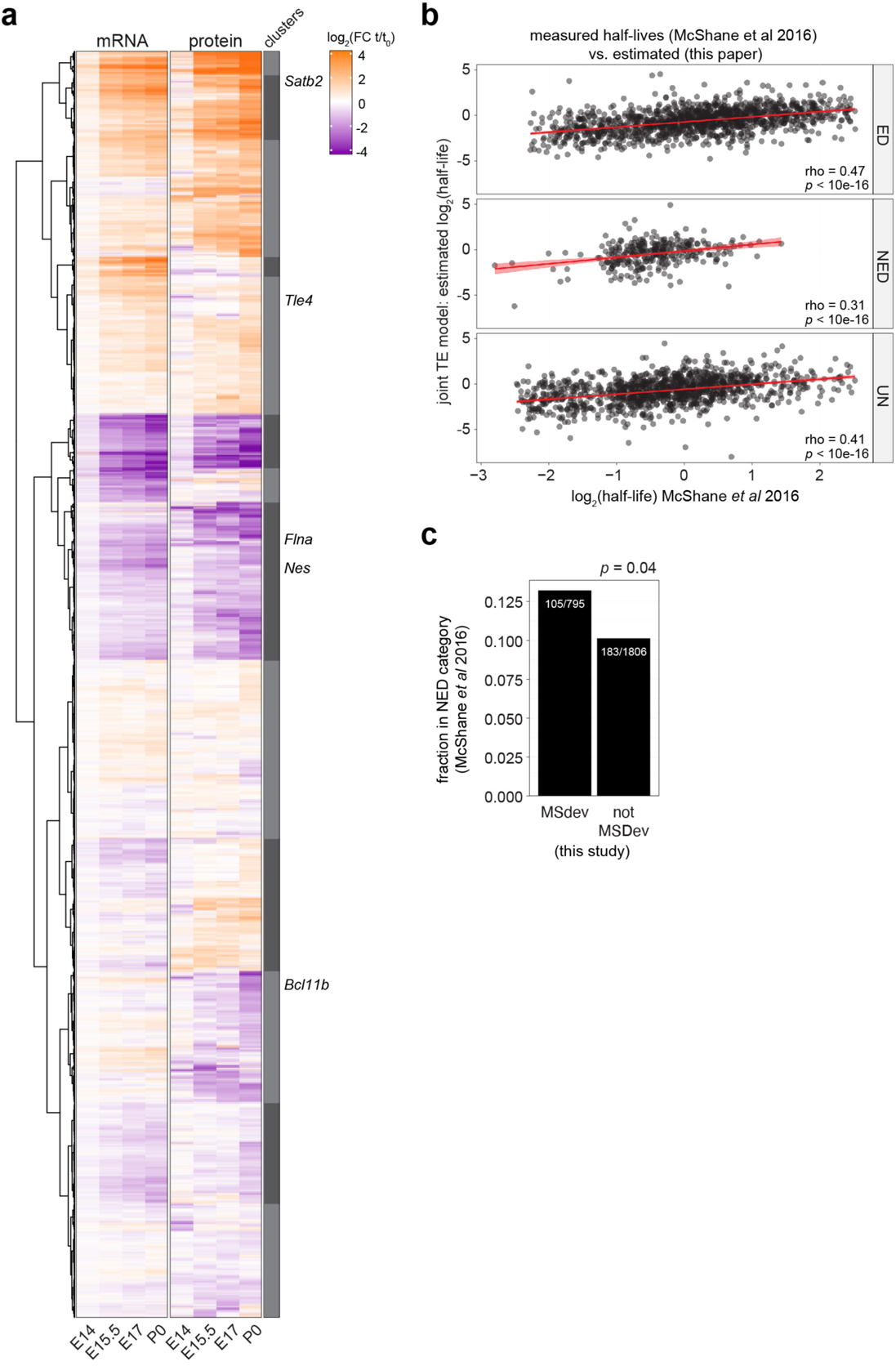
Modeling of mRNA translation. **a**, Hierarchical clustering based on mRNA (RNA-seq) and protein (MS) expression trajectories per gene. Fold change expression increasing or decreasing from E12.5 (t_0_) to subsequent developmental stages (t) shown in heat map. Neural stem cell and neuronal marker genes are indicated (right). **b**, Protein half-lives measured by SILAC MS and categorized as exponential decay (ED), non-exponential decay (NED), or neither (UN) in ^47^ correlated with the model estimates from our data as per ^46^. **c**, The fraction of genes modeled as MS deviating or non-deviating in this study that are categorized as NED proteins in ^47^. Associated with Fig. 7. See also **Supplementary Table 5**.

**Extended Data Fig. 8.**
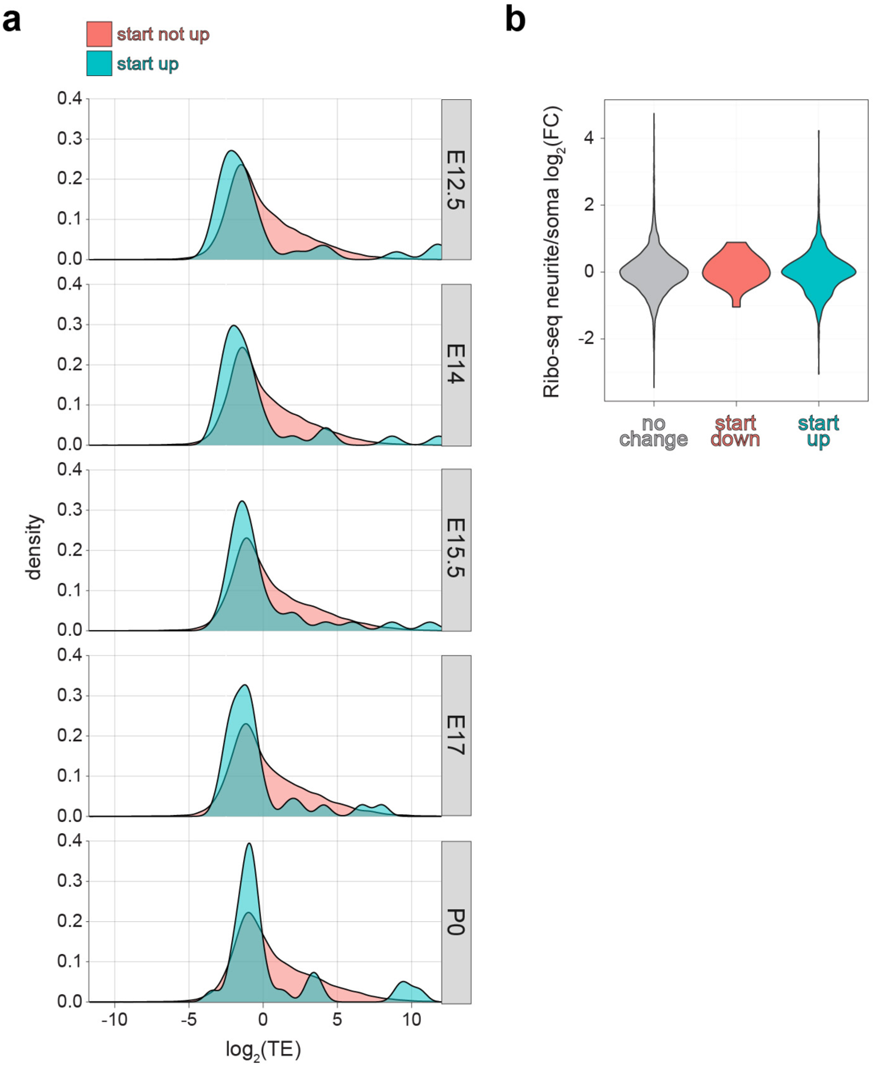
Start codon effect analysis. **a**, TE distribution for genes with increasing start codon occupancy across developmental stages, compared to those without start occupancy changes. **b**, The association of mRNAs demonstrating start codon occupancy changes with translation in neurites vs. the soma of cultured neurons ^81^. Associated with Fig. 5.

## References

1. Buccitelli, C. & Selbach, M. mRNAs, proteins and the emerging principles of gene expression control. Nat. Rev. Genet. 21, 630–644 (2020).

2. DeBoer, E. M., Kraushar, M. L., Hart, R. P. & Rasin, M. R. Post-transcriptional regulatory elements and spatiotemporal specification of neocortical stem cells and projection neurons. Neuroscience 248, 499–528 (2013).

3. Cadwell, C. R., Bhaduri, A., Mostajo-Radji, M. A., Keefe, M. G. & Nowakowski, T. J. Development and Arealization of the Cerebral Cortex. Neuron 103, 980–1004 (2019).

4. Telley, L. et al. Temporal patterning of apical progenitors and their daughter neurons in the developing neocortex. Science. 364, (2019).

5. Zahr, S. K. et al. A Translational Repression Complex in Developing Mammalian Neural Stem Cells that Regulates Neuronal Specification. Neuron 97, 520–537 (2018).

6. Nowakowski, T. J. et al. Spatiotemporal gene expression trajectories reveal developmental hierarchies of the human cortex. Science. 358, 1318–1323 (2017).

7. Klingler, E. et al. Temporal controls over inter-areal cortical projection neuron fate diversity. Nature 599, 453–457 (2021).

8. Llorca, A. et al. A stochastic framework of neurogenesis underlies the assembly of neocortical cytoarchitecture. Elife 8, e51381 (2019).

9. Klingler, E. & Jabaudon, D. Do progenitors play dice? Elife 9, e54042 (2020).

10. Hoye, M. L. & Silver, D. L. Decoding mixed messages in the developing cortex: translational regulation of neural progenitor fate. Curr. Opin. Neurobiol. 66, 93–102 (2021).

11. Kraushar, M. L., Popovitchenko, T., Volk, N. L. & Rasin, M.-R. The frontier of RNA metamorphosis and ribosome signature in neocortical development. Int. J. Dev. Neurosci. 55, 131–139 (2016).

12. Kraushar, M. L. et al. Thalamic WNT3 Secretion Spatiotemporally Regulates the Neocortical Ribosome Signature and mRNA Translation to Specify Neocortical Cell Subtypes. J. Neurosci. 35, 10911–10926 (2015).

13. Kraushar, M. L. et al. Temporally defined neocortical translation and polysome assembly are determined by the RNA-binding protein Hu antigen R. Proc. Natl. Acad. Sci. USA 111, E3815–24 (2014).

14. Zahr, S. K., Kaplan, D. R. & Miller, F. D. Translating neural stem cells to neurons in the mammalian brain. Cell Death Differ. 26, 2495–2512 (2019).

15. Kraushar, M. L. et al. Protein Synthesis in the Developing Neocortex at Near-Atomic Resolution Reveals Ebp1-Mediated Neuronal Proteostasis at the 60S Tunnel Exit. Mol. Cell 81, 304–322.e16 (2021).

16. Popovitchenko, T. et al. Translational derepression of Elavl4 isoforms at their alternative 5′ UTRs determines neuronal development. Nat. Commun. 11, 1674 (2020).

17. Kalish, B. T. et al. Maternal immune activation in mice disrupts proteostasis in the fetal brain. Nat. Neurosci. 24, 204–13 (2021).

18. Ingolia, N. T., Ghaemmaghami, S., Newman, J. R. S. & Weissman, J. S. Genome-wide analysis in vivo of translation with nucleotide resolution using ribosome profiling. Science. 324, 218–223 (2009).

19. Britanova, O. et al. Satb2 Is a Postmitotic Determinant for Upper-Layer Neuron Specification in the Neocortex. Neuron 57, 378–392 (2008).

20. Thoreen, C. C. et al. A unifying model for mTORC1-mediated regulation of mRNA translation. Nature 485, 109–13 (2012).

21. Jin, H. et al. TRIBE editing reveals specific mRNA targets of eIF4E-BP in Drosophila and in mammals. Sci. Adv. 6, eabb8771 (2020).

22. Greig, L. C., Woodworth, M. B., Galazo, M. J., Padmanabhan, H. & Macklis, J. D. Molecular logic of neocortical projection neuron specification, development and diversity. Nat. Rev. Neurosci. 14, 755–69 (2013).

23. Ingolia, N. T. Ribosome Footprint Profiling of Translation throughout the Genome. Cell 165, 22–33 (2016).

24. Dunn, J. G., Foo, C. K., Belletier, N. G., Gavis, E. R. & Weissman, J. S. Ribosome profiling reveals pervasive and regulated stop codon readthrough in Drosophila melanogaster. Elife 2, 1–32 (2013).

25. Li, J. J., Bickel, P. J. & Biggin, M. D. System wide analyses have underestimated protein abundances and the importance of transcription in mammals. PeerJ 2, e270 (2014).

26. Jovanovic, M. et al. Dynamic profiling of the protein life cycle in response to pathogens. Science. 347, 1259038 (2015).

27. Arlotta, P. et al. Neuronal subtype-specific genes that control corticospinal motor neuron development in vivo. Neuron 45, 207–21 (2005).

28. Alcamo, E. a. et al. Satb2 Regulates Callosal Projection Neuron Identity in the Developing Cerebral Cortex. Neuron 57, 364–377 (2008).

29. Frederikson, K. & McKay, R. D. G. Proliferation and differentiation of rat neuroepithelial precursor cells in vivo. J. Neurosci. 8, 1144–1151 (1988).

30. Josephson, R. et al. POU transcription factors control expression of CNS stem cell-specific genes. Development 125, 3087–3100 (1998).

31. Ambrozkiewicz, M. C., Bessa, P., Salazar-Lázaro, A., Salina, V. & Tarabykin, V. Satb2 Cre/+ mouse as a tool to investigate cell fate determination in the developing neocortex. J. Neurosci. Methods 291, 113–121 (2017).

32. Mills, E. W. & Green, R. Ribosomopathies: There’s strength in numbers. Science. 358, eaan2755 (2017).

33. Shah, P., Ding, Y., Niemczyk, M., Kudla, G. & Plotkin, J. B. Rate-limiting steps in yeast protein translation. Cell 153, 1589–1601 (2013).

34. Li, K., Hope, C. M., Wang, X. A. & Wang, J. P. RiboDiPA: a novel tool for differential pattern analysis in Ribo-seq data. Nucleic Acids Res. 48, 12016–12029 (2020).

35. O’Connor, P. B. F., Andreev, D. E. & Baranov, P. V. Comparative survey of the relative impact of mRNA features on local ribosome profiling read density. Nat. Commun. 7, 12915 (2016).

36. Gobet, C. et al. Robust landscapes of ribosome dwell times and aminoacyl-tRNAs in response to nutrient stress in liver. Proc. Natl. Acad. Sci. U. S. A. 117, 9630–9641 (2020).

37. Fang, H. et al. Scikit-ribo Enables Accurate Estimation and Robust Modeling of Translation Dynamics at Codon Resolution. Cell Syst. 6, 180–191.e4 (2018).

38. Riba, A. et al. Protein synthesis rates and ribosome occupancies reveal determinants of translation elongation rates. Proc. Natl. Acad. Sci. 116, 15023–15032 (2019).

39. Weinberg, D. E. et al. Improved Ribosome-Footprint and mRNA Measurements Provide Insights into Dynamics and Regulation of Yeast Translation. Cell Rep. 14, 1787–1799 (2016).

40. Ingolia, N. T., Lareau, L. F. & Weissman, J. S. Ribosome profiling of mouse embryonic stem cells reveals the complexity and dynamics of mammalian proteomes. Cell 147, 789–802 (2011).

41. dos Reis, M., Savva, R. & Wernisch, L. Solving the riddle of codon usage preferences: A test for translational selection. Nucleic Acids Res. 32, 5036–5044 (2004).

42. Chadani, Y. et al. Intrinsic Ribosome Destabilization Underlies Translation and Provides an Organism with a Strategy of Environmental Sensing. Mol. Cell 68, 528–539 (2017).

43. Quax, T. E. F., Claassens, N. J., Söll, D. & van der Oost, J. Codon Bias as a Means to Fine-Tune Gene Expression. Mol. Cell 59, 149–161 (2015).

44. Lennox, A. L., Mao, H. & Silver, D. L. RNA on the brain: emerging layers of post-transcriptional regulation in cerebral cortex development. *WIREs Dev*. Biol. 7, e290 (2018).

45. Schwanhäusser, B. et al. Global quantification of mammalian gene expression control. Nature 473, 337–42 (2011).

46. Becker, K. et al. Quantifying post-transcriptional regulation in the development of Drosophila melanogaster. Nat. Commun. 9, 4970 (2018).

47. McShane, E. et al. Kinetic Analysis of Protein Stability Reveals Age-Dependent Degradation. Cell 167, 803–815.e21 (2016).

48. Schwanhäusser, B., Gossen, M., Dittmar, G. & Selbach, M. Global analysis of cellular protein translation by pulsed SILAC. Proteomics 9, 205–209 (2009).

49. Pechmann, S. & Frydman, J. Evolutionary conservation of codon optimality reveals hidden signatures of cotranslational folding. Nat. Struct. Mol. Biol. 20, 237–243 (2013).

50. Heyer, E. E. & Moore, M. J. Redefining the Translational Status of 80S Monosomes. Cell 164, 757–769 (2016).

51. Verma, M. et al. A short translational ramp determines the efficiency of protein synthesis. Nat. Commun. 10, 5774 (2019).

52. Gingold, H. et al. A dual program for translation regulation in cellular proliferation and differentiation. Cell 158, 1281–1292 (2014).

53. Ishimura, R. et al. Ribosome stalling induced by mutation of a CNS-specific tRNA causes neurodegeneration. Science. 345, 455–9 (2014).

54. Vaninsberghe, M., Berg, J., Andersson-rolf, A., Clevers, H. & Oudenaarden, A. Single-cell Ribo-seq reveals cell cycle-dependent translational pausing. Nature 597, 561–565 (2021).

55. Slavov, N. Unpicking the proteome in single cells. Science. 367, 512–513 (2020).

56. Brunner, A.-D. et al. Ultra-high sensitivity mass spectrometry quantifies single-cell proteome changes upon perturbation. bioRxiv (2020). doi:10.1101/2020.12.22.423933

57. Ambrozkiewicz, M. C. & Kawabe, H. HECT-type E3 ubiquitin ligases in nerve cell development and synapse physiology. FEBS Lett. 589, 1635–1643 (2015).

58. Chiang, S. Y. et al. Usp11 controls cortical neurogenesis and neuronal migration through Sox11 stabilization. Sci. Adv. 7, eabc6093 (2021).

59. Ambrozkiewicz, M. C. et al. Polarity Acquisition in Cortical Neurons Is Driven by Synergistic Action of Sox9-Regulated Wwp1 and Wwp2 E3 Ubiquitin Ligases and Intronic miR-140. Neuron 100, 1097–1115.e15 (2018).

60. Wang, X., Park, J., Susztak, K., Zhang, N. R. & Li, M. Bulk tissue cell type deconvolution with multi-subject single-cell expression reference. Nat. Commun. 10, (2019).

61. Jew, B. et al. Accurate estimation of cell composition in bulk expression through robust integration of single-cell information. Nat. Commun. 11, 1971 (2020).

62. Harris, B. D., Crow, M., Fischer, S. & Gillis, J. Single-cell co-expression analysis reveals that transcriptional modules are shared across cell types in the brain. Cell Syst. 12, 748–756.e3 (2021).

63. Martin, M. Cutadapt removes adapter sequences from high-throughput sequencing reads. EMBnet.journal 17, 10–11 (2011).

64. Langmead, B. & Salzberg, S. L. Fast gapped-read alignment with Bowtie 2. Nat. Methods 9, 357–359 (2012).

65. Dobin, A. et al. STAR: ultrafast universal RNA-seq aligner. Bioinformatics 29, 15–21 (2013).

66. Law, C. W., Chen, Y., Shi, W. & Smyth, G. K. Voom: Precision weights unlock linear model analysis tools for RNA-seq read counts. Genome Biol. 15, R29 (2014).

67. Xiao, Z., Zou, Q., Liu, Y. & Yang, X. Genome-wide assessment of differential translations with ribosome profiling data. Nat. Commun. 7, 1–11 (2016).

68. Calviello, L., Sydow, D., Harnett, D. & Ohler, U. Ribo-seQC: comprehensive analysis of cytoplasmic and organellar ribosome profiling data. bioRxiv (2019). doi:10.1101/601468

69. Ahmed, N. et al. Identifying A- and P-site locations on ribosome-protected mRNA fragments using Integer Programming. Sci. Rep. 9, 6256 (2019).

70. Cui, H., Hu, H., Zeng, J. & Chen, T. DeepShape: Estimating isoform-level ribosome abundance and distribution with Ribo-seq data. BMC Bioinformatics 20, 678 (2019).

71. Patro, R., Duggal, G., Love, M. I., Irizarry, R. A. & Kingsford, C. Salmon provides fast and bias-aware quantification of transcript expression. Nat. Methods 14, 417–419 (2017).

72. Köster, J. & Rahmann, S. Snakemake - a scalable bioinformatics workflow engine. Bioinformatics 28, 2520–2522 (2012).

73. Soneson, C., Love, M. I. & Robinson, M. D. Differential analyses for RNA-seq: Transcript-level estimates improve gene-level inferences. F1000Research 4, 1–19 (2016).

74. Cox, J. & Mann, M. MaxQuant enables high peptide identification rates, individualized p.p.b.-range mass accuracies and proteome-wide protein quantification. Nat. Biotechnol. 26, 1367–1372 (2008).

75. Cox, J. et al. Andromeda: A peptide search engine integrated into the MaxQuant environment. J. Proteome Res. 10, 1794–1805 (2011).

76. Cox, J. et al. Accurate Proteome-wide Label-free Quantification by Delayed Normalization and Maximal Peptide Ratio Extraction, Termed MaxLFQ. Mol. Cell. Proteomics 13, 2513–2526 (2014).

77. McLeay, R. C. & Bailey, T. L. Motif Enrichment Analysis: A unified framework and an evaluation on ChIP data. BMC Bioinformatics 11, 165 (2010).

78. Ray, D. et al. A compendium of RNA-binding motifs for decoding gene regulation. Nature 499, 172–177 (2013).

79. Schindelin, J. et al. Fiji: An open-source platform for biological-image analysis. Nat. Methods 9, 676–682 (2012).

80. Ambrozkiewicz, M. C. et al. The murine ortholog of Kaufman oculocerebrofacial syndrome protein Ube3b regulates synapse number by ubiquitinating Ppp3cc. Mol. Psychiatry 26, 1980–1995 (2020).

81. Zappulo, A. et al. RNA localization is a key determinant of neurite-enriched proteome. Nat. Commun. 8, 583 (2017).

